# GLE1 dysfunction compromises cellular homeostasis, spatial organization, and peripheral axon branching

**DOI:** 10.1101/2024.10.14.618161

**Authors:** T Zárybnický, S Lindfors, S Metso, Z Szabo, R Valtonen, M Tulppo, J Magga, S Saarimäki, S Bläuer, I Miinalainen, R Kerkelä, J Väänänen, R Kivelä, FP Zhang, P Sipilä, R Hinttala, S Kuure

## Abstract

The GLE1 protein is an enigmatic factor of RNA processing, associated with multiple developmental disorders including lethal congenital contracture syndrome 1 (LCCS1). Using *in vivo* genetic engineering to study disturbed GLE1 functions under physiological conditions we demonstrate that inactivation of *Gle1* impedes cellular function and organization and causes pre-gastrulation lethality due to defects in adhesion and lineage specification. In contrast, the knock-in mice genocopying LCCS1-associated *GLE1*_FinMajor_ variant (*Gle1*^PFQ/PFQ^) survive prenatal period but die suddenly at mid-adulthood. *Gle1*^PFQ/PFQ^ mice present irregular count and distribution of spinal motor neurons and impaired development of neural crest-derived tissues as demonstrated by defects in their sympathetic innervation of heart ventricles, paravertebral sympathetic ganglia volume, and adrenal chromaffin cell counts. Unlike previously reported for yeast and HeLa cells, analysis of molecular consequences of *GLE1*_FinMajor_ variant identified normal poly(A)+ RNA distribution in *Gle1*^PFQ/PFQ^ cells, which however were impaired in RNA and protein synthesis and simultaneously showed typical signs of cellular senescence. *Gle1*^PFQ/PFQ^ also induced disturbed stress responses with significant changes in G3BP1-positive stress granule count. Our results show necessity of GLE1 functions for life and indicate that LCCS1 etiology is resultant of pathogenic *GLE1*_FinMajor_ variant impinging differentiation of neural crest derivatives and leading to complex multiorgan defects.

**Highlights:**

- Total inactivation of GLE1 results in disorganization of blastocyst inner cell mass and early embryonic lethality.
- The *Gle1* knock-in (KI) mice, which genocopy the human *GLE1*_FinMajor_ variant causative for lethal congenital contracture syndrome 1 (LCCS1), die suddenly in mid-adulthood.
- Normal poly(A)+ RNA distribution was observed in *Gle1* KI cells, but decreased number of G3BP1-positive stress granules were detected in response to stress.
- Abnormal sympathetic innervation of heart ventricles was detected in *Gle1* KI mice.
- Neural crest-derived tissues represent a new target of *GLE1*_FinMajor_ and GLE1-related disorders.

## Introduction

The fate of an mRNA transcript is determined by stepwise changes in messenger ribonucleoprotein complexes, achieved through the remodeling actions of RNA-dependent DEAD-box ATPases (DDX in eukaryotic cells, Dbp in *Saccharomyces cerevisiae*). Providing an additional layer of regulation, some Dbps/DDXs require specific cofactors to modulate their ATPase activities. Among the known cofactors, the conserved multidomain protein GLE1 is a uniquely versatile DDX/Dbp modulator^1^ playing a direct role in poly(A)+ RNA nucleocytoplasmic shuttling^2^, protein translation^3^, and formation of cytosolic stress granules^3^.

GLE1 dysfunction has been associated with several devastating diseases including ALS^4^, rare diseases enriched in isolated populations like lethal congenital contracture syndrome 1^5^ (LCCS1, OMIM #253310, autosomal recessive), and congenital arthrogryposis with anterior horn cell disease (CAAHD, OMIM #611890, autosomal recessive-homozygous or compound heterozygous mutation), as well as other developmental disorders^6–8^. GLE1 malfunctions have so far been studied *in vivo* in zebrafish (*Danio rerio*), where its deficiency demonstrated a complex phenotype including immobility, edema, underdeveloped motor neurons, apoptosis in the central nervous system^9^, and defective myelination of Schwann cells^10^. The changes partially mimic lethal congenital contracture syndrome 1 (LCCS1), which is an autosomal recessive disease causally linked to the GLE1 dysfunction^11^. LCCS1 is typically observed in early pregnancy due to fetal akinesia and it invariably leads to prenatal death of the fetus before the 32nd gestational week^5,12^. The cause of death is obscure, but the fetuses show hydrops, pulmonary and skeletal muscle hypoplasia, micrognathia and arthrogryposis, defects in anterior horn motor neurons, and severe atrophy of the ventral spinal cord. The most common variant associated with LCCS1 is *GLE1*_FinMajor_ (c.432-10A>G), which is significantly enriched in ethnic Finns. The aberration results in alternative splicing of the *GLE1* transcript inducing an intronic retention of nine nucleotides, and thus coding extra three amino acids (PFQ) in the mutant GLE1 protein.

The *GLE1*_FinMajor_ mutation has been intensely studied *in vitro* in yeast and HeLa cells where its function was associated with disturbed mRNA nucleocytoplasmic shuttling and poly(A)+ RNA accumulation in cell nucleus, as well as malformations and disorganization of GLE1 associated complexes^13^. The *in vivo* consequences of GLE1 dysfunctions in mammals and particularly in LCCS1 and CAAHD pathologies are largely missing. This hampers our understanding of why certain cell populations are more sensitive to RNA dysregulation than others. To elucidate this, we employed mice as a model to genetically disturb GLE1 functions with CRISPR/Cas9 genome engineering. Phenotypic analyses of the two novel mouse models generated, *Gle1* knockout and *Gle1* knock-in of the *GLE1*_FinMajor_ mutation, demonstrate the requirement of GLE1 function for gastrulation and proper development of peripheral nervous system. These findings provide novel aspects on the function of GLE1 in development, health, and disease.

## Results

### Loss of GLE1 function compromises lineage specification in early mouse development

To start elucidating GLE1 physiological functions *in vivo*, we first inactivated the *Gle1* function by utilizing CRISPR/Cas9 system in mouse zygotes. The first exon was targeted by two gRNAs (Figure 1A), leading to CAS9-mediated cut, introduction of an approximately 140 bp deletion (Figures 1B, S1A) and resulting in frameshift mutation corresponding to the premature stop codon (Figure 1C). Heterozygote *Gle1* mice (*Gle1^+/-^*) were born with expected Mendelian ratios, but no homozygote *Gle1* KO mice (*Gle1*^-/-^) were detected after birth (Figure 1D). Systematic stage-by-stage analysis of embryos collected from heterozygote mating revealed the presence of *Gle1*^-/-^ embryos only at embryonic day 3.5 (E3.5) or earlier (Figure 1D). Thus, inactivation of GLE1 function results in much earlier embryonic lethality than that reported for human LCCS1 fetuses^5,12^.

**Figure 1:**
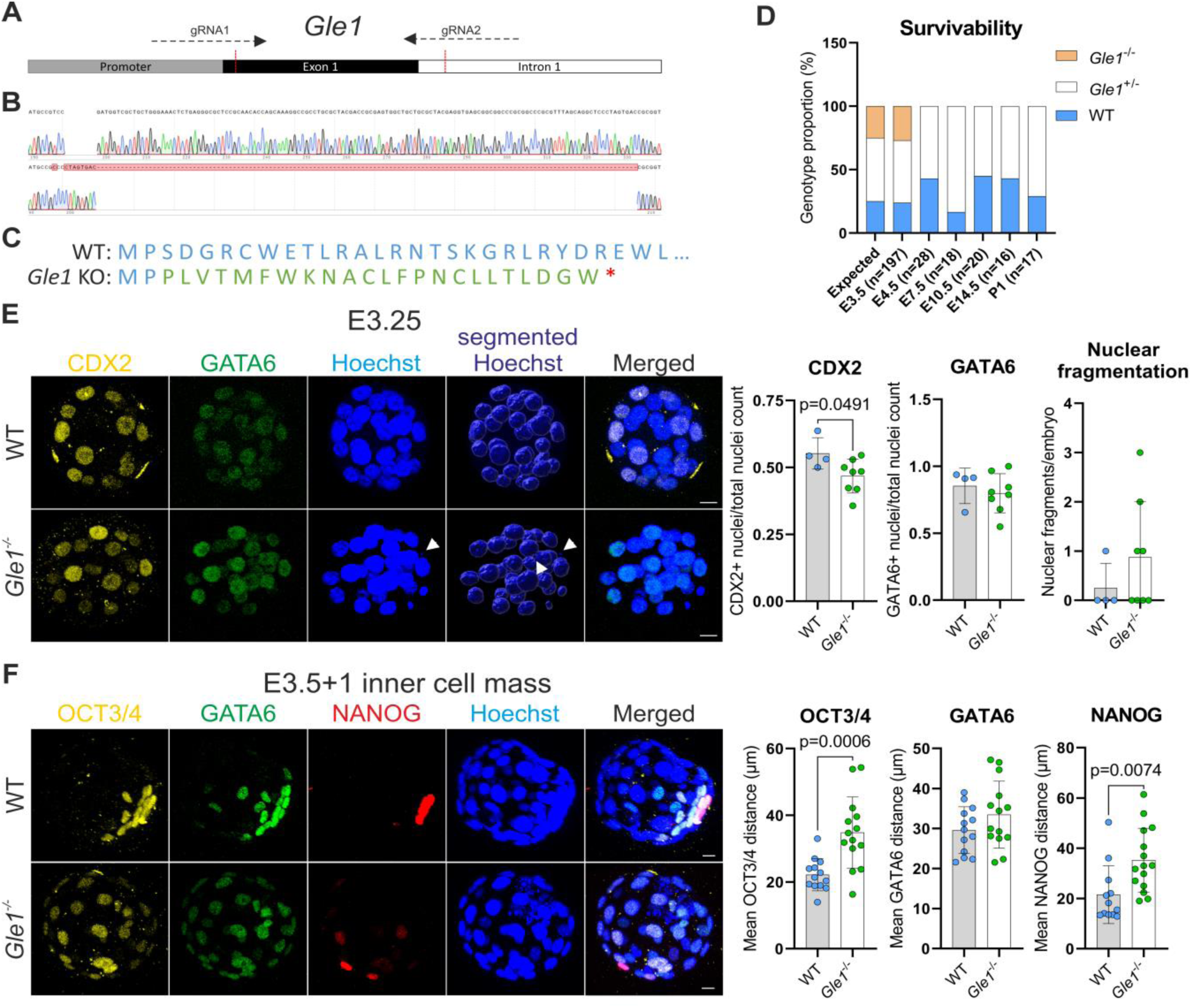
Generation of the *Gle1* KO mice and early embryo phenotype characterization. **A)** Schematics of the strategy for CRISPR/Cas9 mediated genome engineering to target exon 1 of the *Gle1* gene by two gRNAs, aiming to induce reading frame shift and early stop codon for deletion of exon 1, potentially resulting in *Gle1* inactivation (knockout, KO). **B)** Sanger sequencing results of the targeted genomic site showing the wild-type (top) and heterozygote (WT/KO, bottom) sequences in F2 generation. The electropherograms demonstrate a formation of around 140 bp indel in the heterozygote DNA sequence. **C)** Native amino acid sequence of the GLE1 protein (blue) in the wild-type (WT) mice (top) and predicted amino acid sequence of the GLE1 protein post CRISPR/Cas9 mediated deletion of exon 1 (bottom), predicted by Benchling, showing reading frame shift (green) and stop codon formation (red star). **D)** Genotype frequencies of embryos produced from *Gle1*^+/-^ heterozygote to heterozygote mating. E: Embryonic day, P: Postnatal day. **E)** Maximum projections of early embryos cultured from 2-cell stage for 2 days until morulae (E3.25), illustrating the presence of immunostained markers for trophectoderm (CDX2, yellow) and primitive endoderm (GATA6, green) in both WT and Gle1^-/-^ homozygotes. The quantification involves determining the number of CDX2-positive and GATA6-positive nuclei relative to the total number of nuclei, which are visualized by Hoechst staininig (blue) as well the as number of nuclear fragments relative to the total number of nuclei. Means ± SD are plotted, as well as individual data points generated from each embryo (WT: n=4, Gle1^-/-^: n=8). Scale bar 10 µm, two-tailed Student’s *t* test. **F)** Maximum projections of blastocysts flushed from super-ovulated females at E3.5 and cultured for one more day (E3.5+1) and analysis of the immunostained inner cell mass. The quantification involves determining the distance between the nuclei expressing the inner cell mass markers: OCT3/44 (yellow), GATA6 (green, endoderm), NANOG (red, pluripotent epiblast); demonstrating an abnormal distribution of these cells all around the Gle1^-/-^ homozygote blastocyst. Hoechst staining (blue) visualizes all cell nuclei. Means ± SD are plotted, as well as individual data points generated from each embryo (WT: n=13, Gle1^-/-^: n=14). Scale bar 10 µm, two-tailed Student’s *t* test.

We next collected embryos from 2-cell to blastocyst stage and either analyzed them directly or cultured *in vitro* to characterize their developmental and differentiation potential. Analysis of E3.25 morulae showed that overall specification to embryo proper, trophectoderm and primitive endoderm lineages occurred similarly in *Gle1*^-/-^ as in wild type (WT) control embryos (Figure 1E). However, a slight decrease in CDX2-positive cells was detected in *Gle1*^-/-^ embryos, which also demonstrated an increase in OCT3/4-positive cells and nuclear fragmentation (Figure 1E, S1B), suggesting possible morphological or functional defects. At late blastocyst stage an abnormal distribution of cell lineages was observed as demonstrated by an increased mean distance between OCT3/4 and NANOG-positive nuclei in the *Gle1*^-/-^ blastocyst (Figure 1F). This indicates defects in morphological changes that segregate inner cell mass from the other lineages of developing embryo.

Next, 5-ethynyl-2’-deoxyuridine (EdU) incorporation assay was employed to assay the proliferative capacity of *Gle1*^-/-^ embryos. This revealed statistically significant decrease in the proportion of EdU-positive nuclei at E3.5 (Figure S2A). To determine whether *Gle1*^-/-^ blastocysts exhibit reduced proliferation due to increased cell death, apoptosis was measured by TUNEL (terminal deoxynucleotidyl transferase dUTP nick end labeling) and cleaved Caspase 3 (c-CASP3) staining. Indeed, TUNEL-positive nuclei were significantly augmented in the *Gle1*^-/-^ blastocysts at E3.5 and an increase in c-CASP3 abundance was detected at late blastocyst stage (E3.5+1; Figure S2B-C). Such changes in viability at late blastocyst stage likely explain the failure to thrive in the absence of GLE1 functions.

### *Gle1***^-/-^** blastocyst show transcriptional changes in cell adhesion, ion channels and transmembrane transport machinery

In search for transcriptional changes that could suggest mechanisms behind defective development in *Gle1*^-/-^ embryos, a bulk RNA-sequencing (RNA-seq) of the E3.5 blastocysts was performed. Initial quality checks demonstrated good segregation and similar distribution of WT and *Gle1*^-/-^ transcripts and identified 213 differentially expressed genes (151 up- and 62 downregulated) with the cutoff of log2 fold change > 1 and adjusted p-value < 0.05 (Figures 2A, S3A-B, Supplementary Data File 1). The most significantly downregulated transcripts in *Gle1*^-/-^ blastocysts (*Kcnv2*, *Flrt3*, *Gfpt2*, *Myo1g*, unannotated BC051665, *Plcg2* and *Zic3*) failed to be detected in all samples by RNA-seq while showing robust counts in control embryos (Supplementary Data File 1). This suggests that GLE1 function in blastocysts is required for their expression.

**Figure 2:**
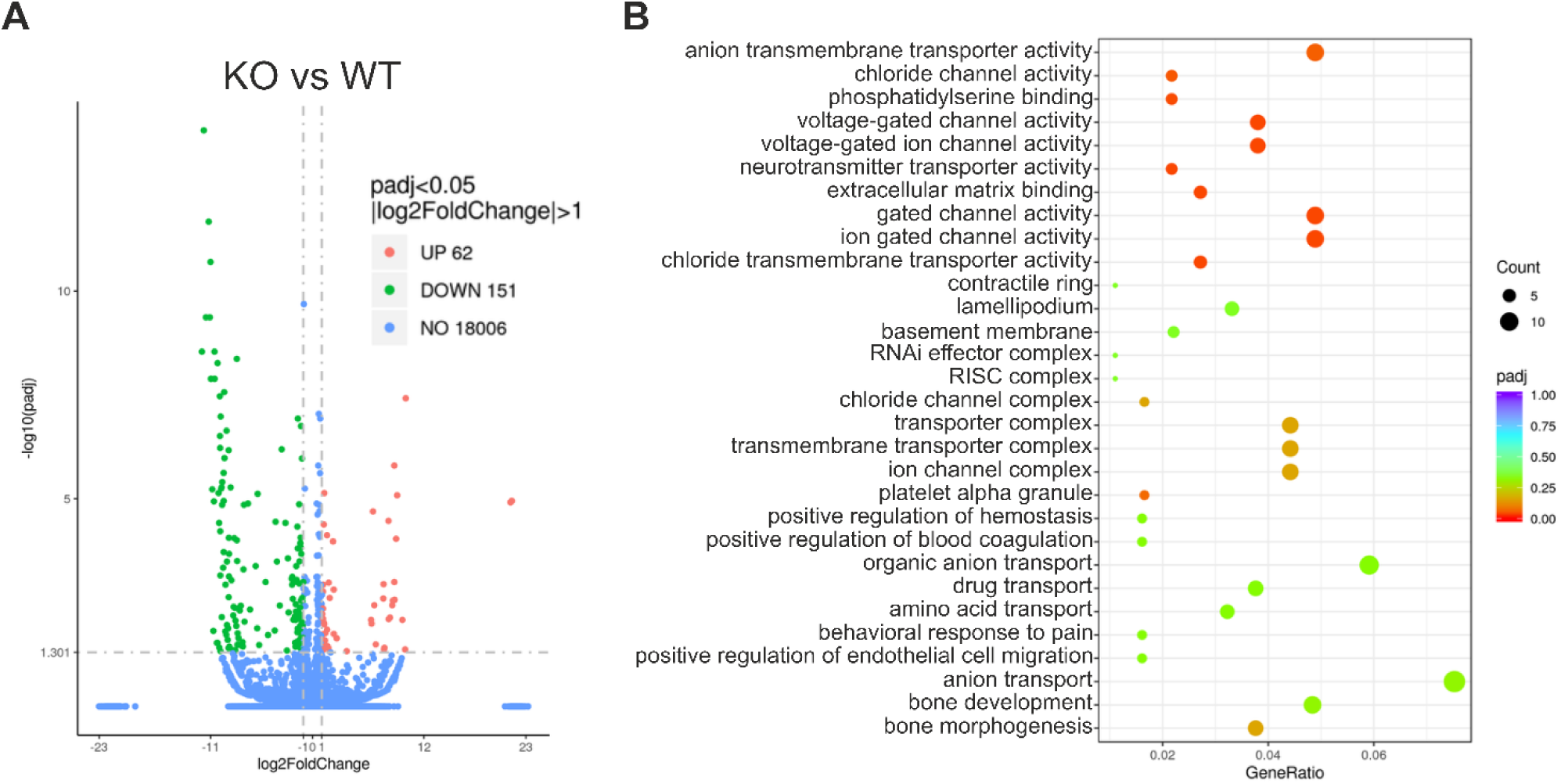
E3.5 *Gle1* KO blastocyst RNA seq. **A)** Volcano plot showing the 213 differentially expressed genes in bulk RNA-seq analysis of E3.5 blastocysts (RNA from 3 blastocysts per biological replicate, three replicates per genotype). Significantly differentially expressed genes in response to *Gle1* knock-out (log2 fold change > 1; adjusted p-value < 0.05) are indicated in green and orange. While 62 genes are upregulated, 151 genes are downregulated. **B)** Dot plot of gene ontology (GO) enrichment analysis visualizing top 30 biological processes affected by *Gle1* loss. Genes with adjusted p-value and log2 fold change > 1 in either direction were used for analysis.

Potassium voltage-gated channel *Kcnv2* (Supplementary Data File 1) encodes the ether-à-go-go-related channel (hERG1) often dysregulated in schizophrenia, cardiac arrhythmia, and tumor proliferation, as well as in normal cell motility involving VIMENTIN (VIM)^14–17^. *Vimentin* (*Vim*) itself and several gene products encoding adhesion and extracellular matrix interacting molecules (*Fltr3*, *Dag1*, *Fndc10*, *Igsf8*, -*9b*, *Cldn23*) and their regulators (*Plcg2*, *Rab15*, *Pigz*, *Ppp6r2*) were among the most significantly downregulated transcripts in *Gle1*^-/-^ embryos (Supplementary Data File 1). Of the most downregulated genes, *Dystroglycan 1* (*Dag1*) inactivation results in early embryonic lethality (E6.5) since it is required for laminin-dependent polarization of epiblast^18,19^. Interestingly, myosin 1G (*Myo1g*) is required for proper cardiac differentiation in *Drosophila*^20^ while loss of zinc finger protein of the cerebellum 3 (*Zic3*) in mice causes neural tube and heart defects^21^.

Among the most significantly upregulated transcripts in *Gle1*^-/-^ embryos were *Cd79b* required for B cell development and *Polh*, which encodes DNA polymerase eta (Supplementary Data File 1) with essential survival functions in replicating damaged DNA^22,23^. Gene ontology (GO) analysis of biological processes revealed the most significant changes in transcripts participating in ion channel and transmembrane transport activities (Figure 2B). This together with altered cellular adhesions suggests that phenotype in early *Gle1*^-/-^ blastocyst development likely derives from defective trans-trophectodermal ion gradient formation, which together with junctional tightening is required for morphological changes^24^ that separate cell lineages physically from each other.

### Knock-in of the LCCS1 *Gle1*^PFQ/PFQ^ variant to mouse genome causes premature death

The early lethality of *Gle1*^-/-^ mice indicates that LCCS1 associated *GLE1*_FinMajor_ variant, which is an intronic mutation, does not fully inactivate GLE1 function, but otherwise modifies its functions. Next, we thus wanted to model LCCS1 in a mammalian model and introduced the human disease causing *GLE1* variant into mouse genome. Although the *GLE1*/*Gle1* exons are highly similar between human and mice, the introns are not. Therefore, it is likely that introducing the intronic A-to-G change in mice would not induce similar alternative splicing as in human patients (Figure 3A). To bypass the species differences in intronic sequences, we straightforwardly knocked in (KI) nine nucleotides (coding the PFQ amino acids) to the intron/exon boundary of mouse *Gle1* gene (Figures 3A, 3B, S4A).

**Figure 3:**
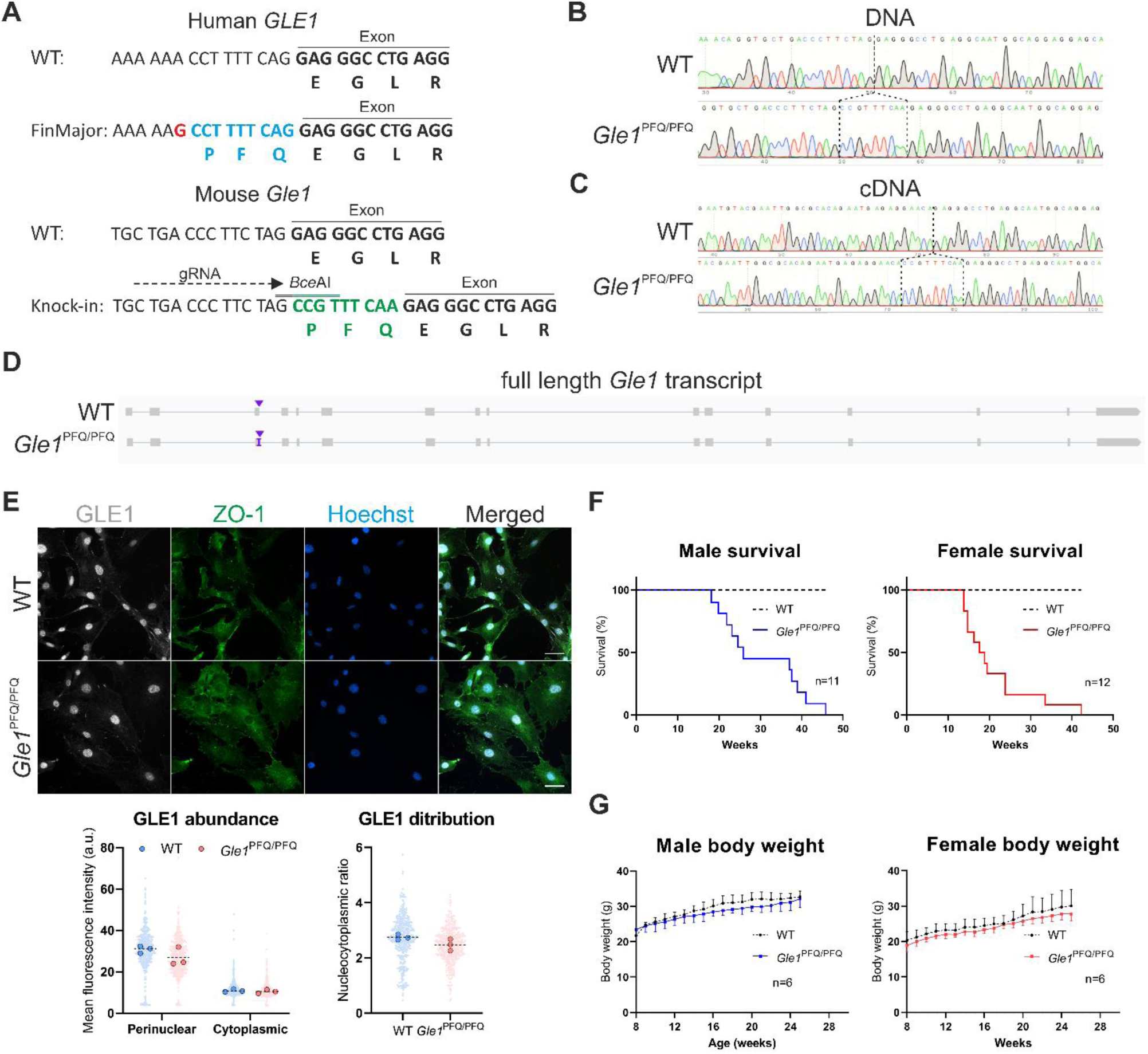
Generation of the *Gle1*^PFQ/PFQ^ KI mice. **A)** Schematics of the target locus comparing the human and mouse sequences of the *GLE1/Gle1* gene, demonstrating similarity in the exon, but major difference in the target intron. This variability mitigates the possibility of introducing the human A-to-G substitution (red), which would induce the alternative splicing and nine nucleotide intron retention (blue), into mice. Nine nucleotides were directly knocked-in (green) at the mouse intron/exon boundary, mimicking the human alternative splicing outcome, with two silent mutations including a digestion site for genotyping purpose (double above line). **B)** Sequence results of the target site from the DNA of wild-type (top) and KI/KI homozygote (bottom) mice, isolated from ear samples of the F2 generation mice. The Sanger sequencing electropherograms demonstrate a successful knock-in of the desired nine nucleotides at the intron/exon boundary. **C)** Sequence results of the target site from the reverse transcribed cDNA of wild-type (top) and KI/KI homozygote (bottom) mice. RNA was isolated from the heart of adult (10 weeks old) mice at the F2 generation. The Sanger sequencing electropherograms demonstrate a successful transcription of the inserted nine nucleotides in the KI homozygotes, mimicking the outcome of the alternative splicing in the GLE1_FinMajor_ mutation. **D)** Analysis of the *Gle1* transcript full length and sequence in the WT and *Gle1* KI mice. Visualization of long-range RNA-seq reads in IGV, produced by the PacBio SMRT sequencing and using the RNA isolated from the postnatal day zero (P0) hearts. **E)** Immunofluorescent analysis of Gle1 (grey) protein abundance and distribution in mouse embryonic fibroblasts isolated at embryonic day 13.5. Individual data points generated from each cell are plotted (150 cells per embryo, three embryos per genotype), as well as median values from each embryo (large dots). Scale bar: 50 µm. **F)** Kaplan-Meier survival curve of the WT and Gle1 KI males and females, showing the premature lethality of KI homozygotes in mid-adulthood. **G)** Analysis of the body weight in time between the adult WT and KI homozygote males and females.

The *Gle1* KI mice were born at Mendelian ratios and successful transcription of the inserted nine nucleotides was confirmed in adult *Gle1*^PFQ/PFQ^ KI homozygotes (Figures 3C, S4B). The result demonstrated a similar transcriptional outcome as detected in cDNA of human *GLE1*_FinMajor_ fetuses^5^. To confirm that the modification did not induce unintentional splicing events of the *Gle1* transcript, we performed full-length transcript sequencing using the PacBio SMRT (Single Molecule, Real-Time). This verified the length and sequence for mouse WT and KI *Gle1* transcripts (Figure 3D) and demonstrated the lack of undesired splicing. No changes in the expression of *Gle1* transcript or protein levels were detected between WT and *Gle1*^PFQ/PFQ^ adult mouse hearts (Figures S4C, S4D), nor did the GLE1 abundance or distribution change in mouse embryonic fibroblasts (MEFs) (Figure 3E). While the LCCS1 patients die during the fetal development, the *Gle1*^PFQ/PFQ^ mice survive until mid-adulthood (Figure 3F) before dying suddenly overnight, without any obvious changes in growth, welfare, or body weight (Figure 3G).

### *Gle1 ^PFQ/PFQ^* KI mouse embryonic fibroblasts demonstrate signals of premature senescence

To dissect the effects of KI modification, we analyzed basic cellular properties in MEFs. Flow cytometry analysis of MEFs showed an increase in the G2/M population of *Gle1*^PFQ/PFQ^ cells (Figure S5A) while the EdU incorporation assay demonstrated a lower rate of proliferation in *Gle1*^PFQ/PFQ^ MEFs than in WT cells (Figure 4A). Interestingly, a striking increase in the number of phospho-histone H3 (Ser10)-positive (pHH3) mitotic nuclei was detected in *Gle1*^PFQ/PFQ^ MEFs (Figures 4A, 4B). This suggests that the *GLE1*_FinMajor_ variant severely disrupts cell cycle progression as *Gle1*^PFQ/PFQ^ MEFs are dominantly mitotic suggesting that they are arrested in M phase. Similar cell cycle progression defect was previously described after the knockdown of *GLE1* in HeLa cells^25^.

**Figure 4:**
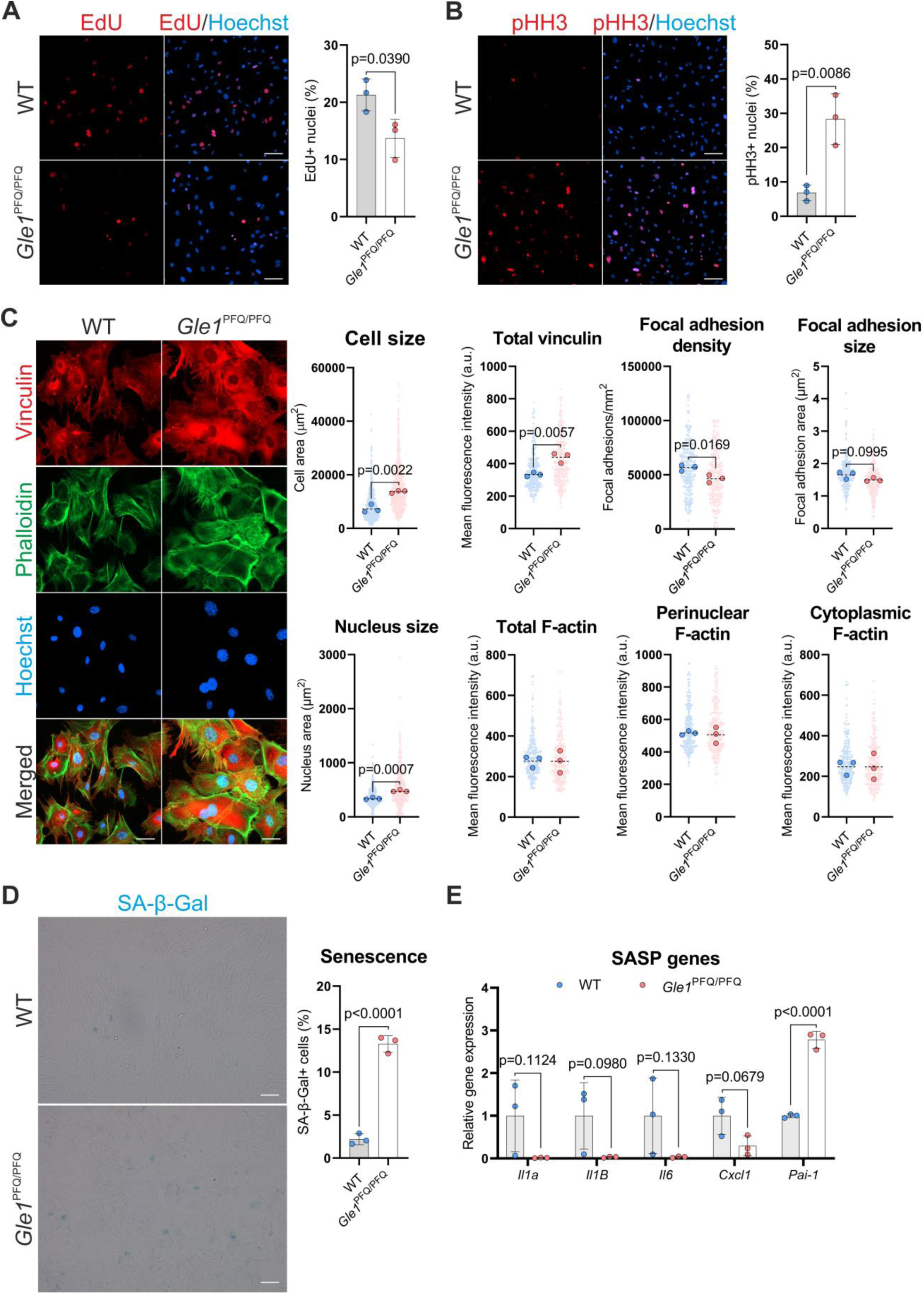
Disturbed cell cycle and senescence progression *Gle1*^PFQ/PFQ^ MEFs. **A)** Cell proliferation assay with quantification of EdU uptake (red, 10 µM EdU, 6 hours). Means ± SD are plotted, as well as individual data points generated from each embryo (three embryos per genotype, over 500 cells analyzed per embryo). Scale bar 100 µm, two-tailed Student’s *t* test. **B)** Immunofluorescent staining and quantification of pHH3+ (red) mitotic cells. Means ± SD are plotted, as well as individual data points generated from each embryo (three embryos per genotype, over 500 cells analyzed per embryo). Scale bar 100 µm, two-tailed Student’s *t* test. **C)** Vinculin and phalloidin stained MEFs were used to analyze the cell size, focal adhesions, and F-actin distribution. Individual data points generated from each cell are plotted (100 cells per embryo, three embryos per genotype), as well as median values from each embryo (large dots). Scale bar 50 µm, two-tailed Student’s *t* test. Hoechst staining (blue) visualizes all cell nuclei in A-C. **D)** SA-β-Gal staining (blue) of MEFs as a marker of cellular senescence. Means ± SD are plotted, as well as individual data points generated from each embryo (three embryos per genotype, over 400 cells analyzed per embryo). Scale bar 200 µm, two-tailed Student’s *t* test. **E)** RT-qPCR quantification of senescence associated secretory profile (SASP) genes in MEFs at passage 4. Means ± SD are plotted, as well as individual data points generated from each embryo (three embryos per genotype). Two-tailed Student’s *t* test.

The cell morphology of *Gle1*^PFQ/PFQ^ MEFs differed by eye from WT cells and prompted us to study this more in detail. Analysis of cell size revealed that *Gle1*^PFQ/PFQ^ cells are significantly larger than wild type cells (Figure 4C). This is shown by the abundance of vinculin-positive focal adhesions, which in *Gle1*^PFQ/PFQ^ cells showed similar size as in wild type cells but significant decrease in the density (Figure 4C). No measurable alterations in cytoskeletal organization as assayed by F-actin distribution and abundancy were detected (Figure 4C).

The significant increase of *Gle1*^PFQ/PFQ^ cells in mitosis together with the observed cytomorphological changes in cell size resemble hallmarks of cellular senescence^26–28^. Such growth arrest is often triggered by persistent response to DNA damage or stress signaling^29–32^, and senescent cells usually exhibit increased lysosomal β-galactosidase activity^33^. Under certain conditions senescent cells also secrete interleukins, inflammatory cytokines, and growth factors which is recognized as the senescent-associated secretory profile (SASP)^34,35^. To further analyze possible premature senescence in *Gle1*^PFQ/PFQ^ MEFs, we utilized senescence-associated β-galactosidase (SA-β-gal) and DNA damage assays. Strikingly, while the wild type cells barely showed any β-gal-positive cells, their abundance in *Gle1*^PFQ/PFQ^ MEFs was significantly increased (Figure 4D). Analysis of DNA damage by staining with phospho-histone H2A.X (Ser 139) (γH2A.X) showed significant increase in the in *Gle1*^PFQ/PFQ^ cells, which additionally showed changes in the expression of cell cycle regulators *Cdkn2a* (p16, p19) and *Cdkn1a* (p21) (Figure S5B-C). Out of the tested SASP related genes, the expression of *Il1α*, *Il1β*, *Il6*, and *Cxcl1* seemed slightly downregulated while *Pai-1* presented a significantly increased expression in the *Gle1*^PFQ/PFQ^ MEFs (Figure 4E). Thus, our characterization of *Gle1*^PFQ/PFQ^ cellular features indicates that genetic LCCS1 variant *Gle1*^PFQ/PFQ^ induces cellular senescence in mouse MEFs.

### *Gle1*^PFQ/PFQ^ variant in mouse allows intact mRNA shuttling but disrupts stress response

Previous *in vitro* experiments have established a basis for *GLE1*_FinMajor_ driven pathology, which lies in disturbed nucleocytoplasmic shuttling of poly(A)+ RNA leading to its accumulation in the cell nucleus^13^. Interested if this is the potential cellular mechanism disrupting GLE1 functions in mouse *Gle1*^PFQ/PFQ^ cells as well, we analyzed the mRNA transport by oligo d(T) fluorescent *in-situ* hybridization (FISH) followed by quantification of poly(A)+ RNA distribution. These experiments found no changes in the oligo d(T) abundance or nucleocytoplasmic distribution between genotypes (Figure 5A). Motivated by the known involvement of GLE1 in translation^3^ we further investigated, if despite normal poly(A)+ RNA distribution, *Gle1*^PFQ/PFQ^ variant would impact RNA and/or protein synthesis. Metabolic labelling through incorporation of ethynyl uridin (EU) indeed identified decrease in RNA synthesis while puromycin incorporation demonstrated diminished protein synthesis, especially in those molecules around the size of 50-75 kDa (Figures S6A, S6B). Notably, these experiments cannot distinguish if the identified changes in RNA synthesis and protein translation are primarily induced by modified GLE1 functions or secondarily by changes in its cellular activity.

**Figure 5:**
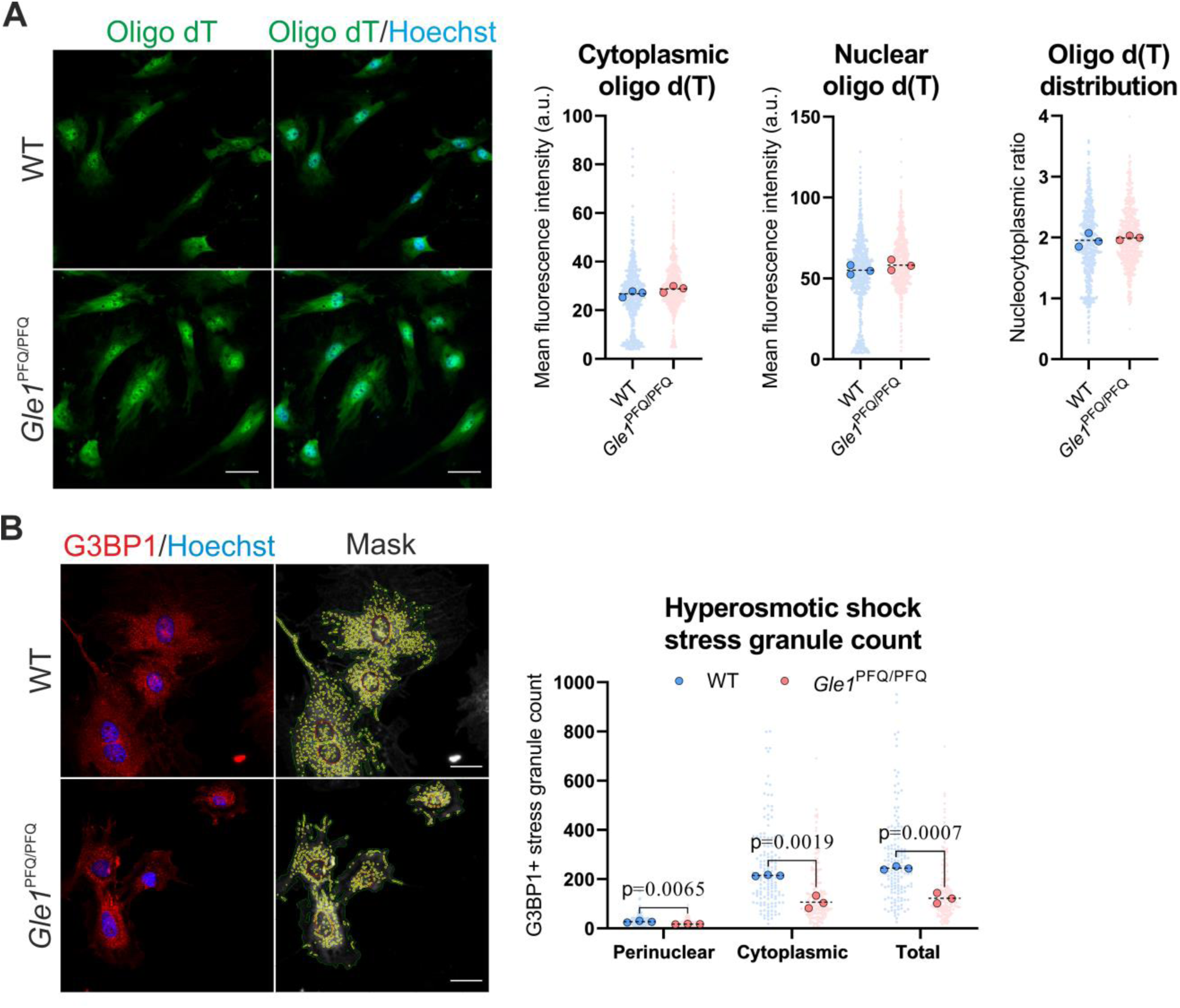
GLE1 functional analysis using E13.5 mouse embryonic fibroblasts. **A)** Visualization and quantification of poly(A)+ RNA through fluorescent *in situ* hybridization of 5’-Cy3-oligo d(T)30 probe in MEFs. Individual data points generated from each cell are plotted (200 cells per embryo, three embryos per genotype), as well as median values from each embryo (large dots). Scale bar 50 µm. **B)** Maximum projection of MEFs stained with G3BP1 (red) after stress challenge with hyperosmotic shock (400mM sorbitol, 1 hour) followed by stress granule quantification as G3BP1-positive speckle type objects (yellow) in the cytoplasm (green) and around the nucleus (red). Hoechst staining (blue) visualizes all cell nuclei. Individual data points generated from each cell are plotted (50 cells per embryo, three embryos per genotype), as well as median values from each embryo (large dots). Scale bar 30 µm, two-tailed Student’s *t* test.

To further assess possible pathological mechanisms beyond mRNA nucleocytoplasmic transport, we next addressed the ability of *Gle1*^PFQ/PFQ^ cells to handle and react to stress, which typically induces integrated stress response (ISR). This pathway is activated by phosphorylation of the alpha subunit of eukaryotic translation initiation factor 2 (eIF2α, serine 51) to tackle a range of physiological changes and several different pathological conditions^36–39^. Activation of ISR modulates the transcriptional and translational responses of the stressed cells to facilitate handling of the immediate effects of the thread^40^.

At sub-cellular level, an acute response to environmental stressors and activation of the ISR promotes stress granule formation^41,42^. We stressed the WT and *Gle1*^PFQ/PFQ^ MEFs with three different conditions: hyperosmotic shock (sorbitol), oxidative shock (sodium arsenite), and heat shock (43°C) and measured p-eIF2α levels to observe similar ISR activation in both genotypes (Figure S5C). However, quantification of stress granule formation by visualizing the core stress granule protein G3BP stress granule assembly factor 1 (G3BP1) demonstrated a significant decrease of the total count of G3BP1-positive stress granules per cell in *Gle1*^PFQ/PFQ^ MEFs under all of tested conditions (Figures 5B, S6D-E). This shows that *Gle1*^PFQ/PFQ^ MEFs have altered capacity to handle stress although their ISR pathway remains intact and suggests essential functions for GLE1 in cellular stress response.

### Impaired motor neuron organization in the developing ventral spinal cord of *Gle1*^PFQ/PFQ^ mice

Following the detailed analysis of morphological and functional changes in *Gle1* KI MEFs, we focused on examining the developmental abnormalities induced by the *Gle1*^PFQ/PFQ^ variant in mouse. One of the most profound phenotypes of the LCCS1 is the loss of ventral horn motor neurons, resulting in fetal akinesia^5^. Primary characterization of E11.5 brachial spinal cords revealed comparable count of OLIG2-positive motor neuron (MN) progenitors in WT and *Gle1* KI embryos (Figure S7A). However, statistically significant reduction in ISL1/2-positive general MNs was detected in the lateral motor column of *Gle1*^PFQ/PFQ^ embryos (Figure S7B) verifying the mouse model phenotype similar but clearly milder than that reported in LCCS1 fetuses. Like in MEFs, oligo d(T) FISH analysis of embryonic spinal cords presented no changes in poly(A) RNA distribution (Figure S7C), supporting our *in vitro* findings that suggest other than global nucleocytoplasmic mRNA shuttling defect as contributing molecular mechanism of LCCS1 pathology.

It has been described that the spinal cord in LCCS1 fetuses is macroscopically thinned because of an early reduction of the anterior horn and a paucity of anterior horn cells^43^. To assess if the *Gle1* KI spinal cords present any morphological phenotype early in the development, we first quantified the number of Sox2-positive progenitors and Sox2/pHH3-positive mitotic cells in the neural tubes without detecting differences between the genotypes (Figure S8A). Next, more detailed scrutiny of ISL1/2-positive general MNs was performed in the E12.5 ventral spinal cord. This revealed revealing that while both brachial and thoracic spinal cord segments presented a lower number of MNs, an increased MN count was surprisingly detected in the lumbar segment (Figure 6A, 6B).

**Figure 6:**
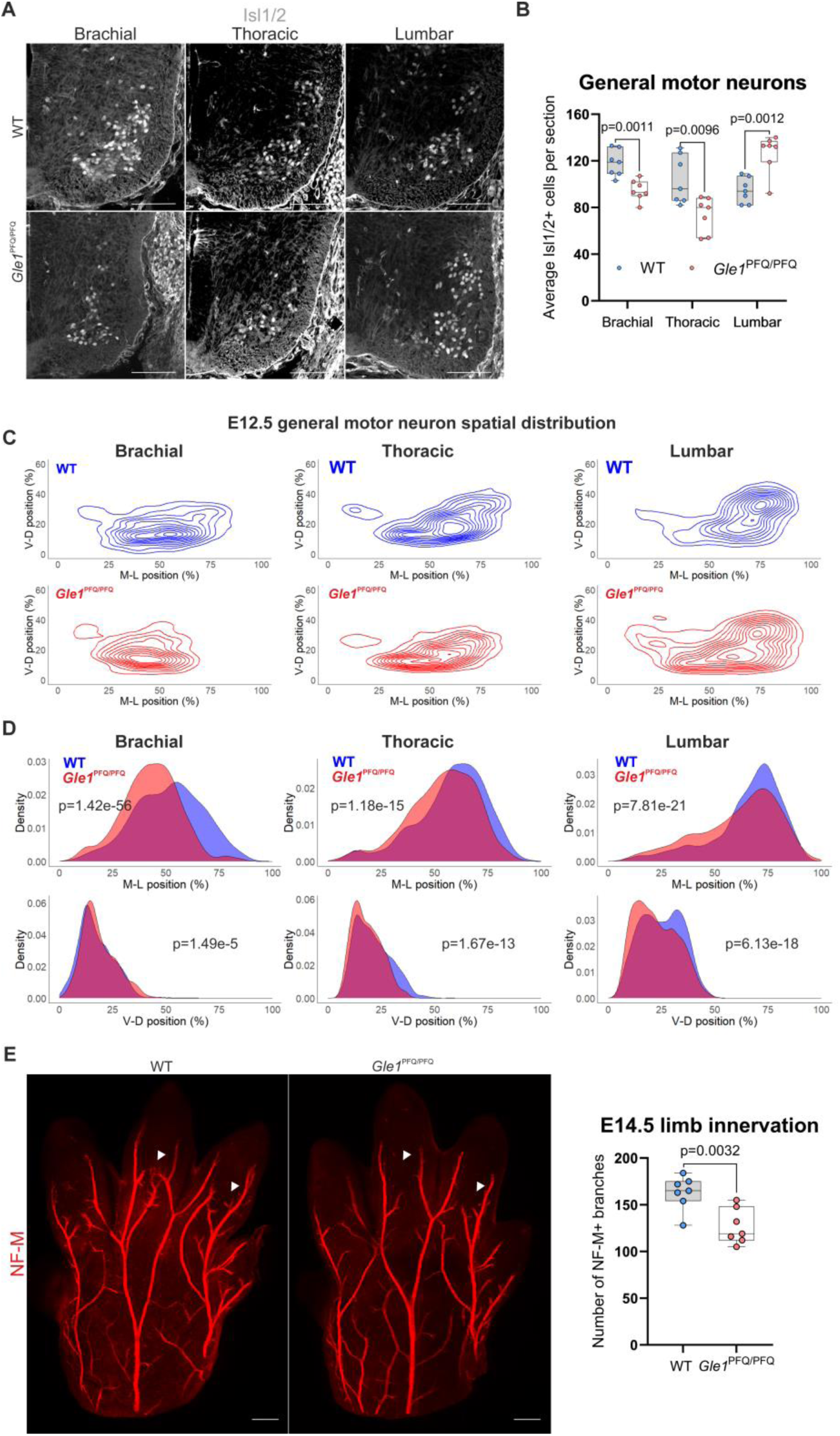
Aberrant number and distribution of spinal cord ventral general motor neurons in E12.5 *Gle1*^PFQ/PFQ^ embryos. **A)** Representative images of ventral spinal cord transverse hemi-sections at the brachial, thoracic, and lumbar regions, immunolabelled for ISL1/2 (white) as a marker of general motor neurons. Scale bar 100 µm. **B)** Quantification of the ISL1/2 immunostained general motor neurons in the ventral spinal cord at the brachial, thoracic, and lumbar regions. Box and whisker plots, as well as individual data points (n=7) generated from each mouse (i.e. the average motor neuron count from multiple sections – brachial: 3, thoracic: 4, lumbar: 4), two-tailed Student’s t test. **C)** Contour density plots demonstrating the qualitative changes in relative spatial distribution of the ventral spinal cord ISL1/2-positive general motor neurons in the medio-lateral (M-D) and ventro-dorsal (V-D) planes. **D)** Density plots highlighting the quantitative changes in relative spatial distribution of the ventral ISL1/2 immunostained general motor neurons in the medio-lateral (M-D) and ventro-dorsal (V-D) planes. Kolmogorov–Smirnov test. **E)** Whole-mount images of neurofilament-M (NF-M, red) immunostained right forelimb paws imaged on the dorsal side, and quantification of the visualized axonal branches. White arrowheads highlight the spots with remarkedly decreased number of branches in the *Gle1*^PFQ/PFQ^ mice. Box and whisker plot, as well as individual data points (n=7) generated from each mouse are shown in the graphs. Scale bar 100 µm

The irregularities in MN numbers are likely to affect the developing neural circuitry and modify the choices for synaptic pruning to optimize the neural network during further development^44^, which can be further vitiated by the shift in MN distribution. Accordingly, the relative localization of MNs in all of the studied spinal cord segments was altered in a way that the MNs in *Gle1*^PFQ/PFQ^ embryos were positioned more medially in the medio-lateral axis than in WT embryos (Figure 6C, 6D). At the ventro-dorsal axis, the *Gle1*^PFQ/PFQ^ brachial MNs were scattered slightly more dorsally, in contrast to the KI thoracic and lumbar MNs shifted more ventrally than the arrangement detected in WTs.

It has been previously described that *gle1* inactivation in zebrafish results not only in fewer motor neurons, but also in disorganized axon arborsation^9^. Encouraged by this we performed whole-mount analysis of E14.5 forelimb paws to address axonal development in *Gle1*^PFQ/PFQ^ mice. This revealed that the main nerve fibers in *Gle1*^PFQ/PFQ^ mice were comparable in structure or organization, but the total number of neurofilament-M-positive axon branches was profoundly decreased *Gle1*^PFQ/PFQ^ mice, mainly due to remarkably diminished axonal side branching (Figure 6E).

### Adrenal chromaffin cells show high prevalence in mitosis but are overall diminished in *Gle1*^PFQ/PFQ^ embryos

Through the pathology observations described in LCCS1 fetuses^45–47^, we hypothesized that neural crest derived tissues might be another affected target structures beyond spinal MNs. To test the hypothesis, we picked the adrenal gland chromaffin cells for screening due to being easily accessible and straightforward to analyze in terms of identifying potential defects. Indeed, a decrease in tyrosine hydroxylase (TH)-positive chromaffin cells and SOX10-positive neural crest cells was detected in *Gle1*^PFQ/PFQ^ embryos in comparison to WT littermates (Figure 7A), supporting our hypothesis. The chromaffin cells retain the ability to proliferate throughout the life^48^ and we became interested in seeing if cell cycle would be disrupted similarly as detected in *Gle1*^PFQ/PFQ^ MEFs. Analysis of pHH3-positive nuclei demonstrated a marked increase, but only in the TH-positive cells of the *Gle1*^PFQ/PFQ^ mutants (Figure 7B), suggesting a cell type-specific sensitivity towards the *Gle1* mutation and a new potential mechanism in the mosaic of LCCS1 and related pathologies.

**Figure 7:**
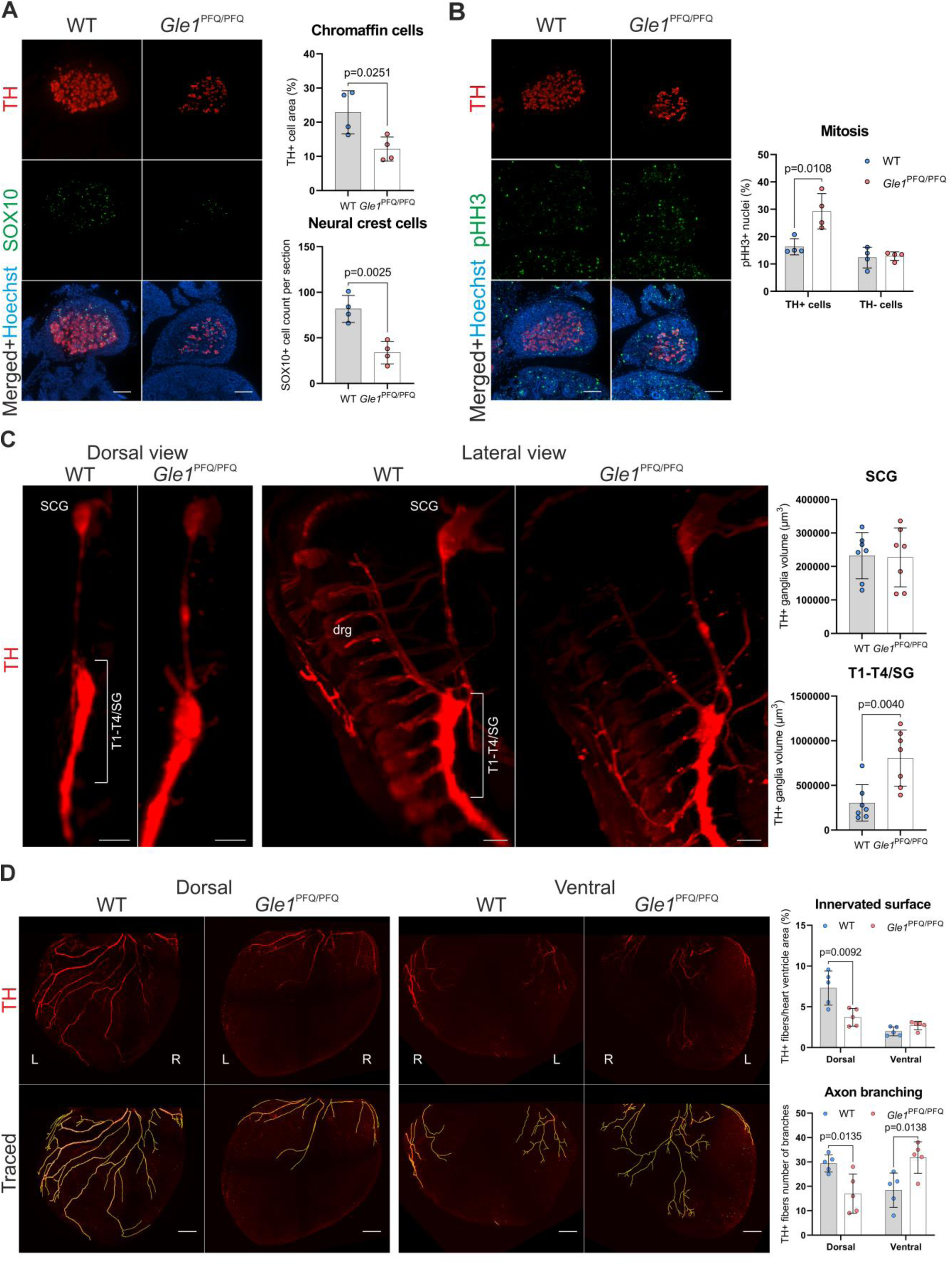
Impaired development of sympathoadrenal system in *Gle1*^PFQ/PFQ^ mouse embryos. **A)** Representative cross-sections of E14.5 adrenal glands immunostained for TH (red, chromaffin cells), and SOX10 (green, neural crest cells). Hoechst staining (blue) visualizes all cell nuclei. Means ± SD are plotted, as well as individual data points generated from each embryo (n=4 per genotype, 3 sections per mouse). Scale bar 100 µm, two-tailed Student’s *t* test. **B)** Representative cross-sections of E14.5 adrenal glands immunostained for TH (red, chromaffin cells), pHH3 (green, mitosis marker) and quantification of mitotic cells in the adrenal gland. Hoechst staining (blue) visualizes all cell nuclei. Means ± SD are plotted, as well as individual data points generated from each embryo (n=4 per genotype, 3 sections per mouse). Scale bar 100 µm, two-tailed Student’s *t* test. **C)** Whole-mount TH immunostaining visualization of the E14.5 paravertebral sympathetic ganglia with focus on the populations contributing to the heart innervation. Quantification of the volume of superior cervical ganglion (SCG), and the block of thoracic T1-T4 and stellate (SG) ganglia (close proximity in early stages, therefore the sympathetic ganglia are distinguished by the position of the nonspecifically stained dorsal root ganglia (drg)). Means ± SD are plotted, as well as individual data points generated from each embryo (n=7 per genotype). Scale bar 50 µm, two-tailed Student’s *t* test. **D)** Whole-mount immunofluorescent staining of tyrosine hydroxylase (TH, red)-positive sympathetic neurons innervating the E16.5 heart ventricles at the dorsal and ventral side. Graphs show quantification of innervated surface area and axon branching. Means ± SD are plotted, as well as individual data points generated from each embryo (n=5 per genotype). Scale bar 200 µm, two-tailed Student’s *t* test.

### GLE1 function is required for normal development of sympathetic nervous system

Excited about the observations in chromaffin cells that are derivatives of the neural crest, but also closely related to sympathetic neurons through sympathoadrenal progenitors^45–47^, we hypothesized that sympathetic nervous system may be similarly affected in the developing *Gle1*^PFQ/PFQ^ mice. We have performed whole-mount imaging of the paravertebral sympathetic ganglia running bilaterally along the spinal cord at the E14.5 stage, which majorly contribute to the sympathetic innervation and control of diverse biological processes, including cardiac output, body temperature, blood glucose levels and immune function under basal conditions and in response to external stressors such as cold or danger^49,50^. Indeed, quantification of TH-positive stellate ganglion and upper thoracic ganglia (SG/T1-T4) volumes revealed significant increase in size in the *Gle1*^PFQ/PFQ^ embryos in comparison to WT controls (Figure 7C), demonstrating an improper developmental process. The SG/T1-T4 block of ganglia majorly contributes towards the sympathetic innervation of heart ventricles^51,52^. Heart is a vital organ supplied with multiple neuronal populations densely innervating the myocardium and which shows high level of neuronal plasticity during injury and disease^53^, and we focused on the E16.5 stage as a critical period of sympathetic neurons populating the heart ventricles. Whole-mount imaging of TH-positive sympathetic neurons in dorsal and ventral side of the ventricles revealed that the innervated surface area in the *Gle1*^PFQ/PFQ^ hearts was significantly smaller on the dorsal side than in wild type embryos. Interestingly, the ventral innervation also demonstrated deviation from the normal arborization by showing much higher number of axonal branches in the *Gle1*^PFQ/PFQ^ than in WT mice (Figure 7D). This disrupted sympathetic patterning of the heart during embryonic development is particularly concerning, as it can lead to postnatal arrhythmias and sudden death^54,55^.

Axon guidance is a complex symphony of intercellular communication, combining attractive and repulsive signals in temporal and spatial order. Considering the unusual pattern of ventricular innervation, we considered other contributing pathological mechanisms and assessed potential changes in cardiac cell communication. Staining E16.5 heart sections with cellular adhesion (N-cadherin), axon guidance (Semaphorin 3A), and cardiomyocyte conductivity (Connexin 43) molecules revealed comparable distribution in left and right ventricles of both genotypes (Figure S9A-C). These results suggest that the abnormal sympathetic ganglia development indeed likely contributes to the aberrant ventricular innervation in our LCCS1 mouse model.

### Adult female *Gle1*^PFQ/PFQ^ mice present signs of heart failure

Thinking of the premature sudden deaths in the KI mice, we next focused on potential long-term consequences of the early cardiac innervation defects in the adult hearts. Analysis of laminin α1 (LAMA1) immunostained cardiomyocytes in 25-week-old hearts identified increased cardiomyocyte size in the left ventricle of the *Gle1*^PFQ/PFQ^ mice (Supplementary figure 10). Interested in if the sympathetic neural network would also be affected in adult hearts, TH-positive sympathetic neurons counterstained with isolectin B4 (IB4) as an endothelial marker were analyzed. This showed a slight decrease in the TH-positive area in the left ventricle, suggesting a minor innervation defect, while the IB4-positive areas presented similar heart vascularization and comparable level of neurovascular interface in both genotypes (Supplementary figure 11A-C).

To evaluate whether the identified changes affect cardiac function, we first performed electrocardiography (ECG) recordings in WT and *Gle1* KI mice at 8 weeks of age to detect no difference in the female mice, whereas male *Gle1* KI mice presented a longer QRS wave duration than WT males (Figure 8A, Supplementary Data File 2). On the other hand, ECG analysis at 16 weeks of age showed no difference in any of the ECG parameters between the genotypes in either females or males (Supplementary Data File 3). To provide insights into the autonomous regulation of heart function, heart rate variability (HRV) analysis was performed from the continuous ECG measurements. This identified an increase in the very low frequency (VLF) bands of unknown etiology in the 8-week-old females, but no other significant changes (Supplementary Data File 4-5). Echocardiography analysis focusing on the systolic function of the left ventricle, which showed increased cardiomyocyte size (Supplementary figure 10), was used next. Analysis of 17–25-week-old mice showed a decrease in stroke volume and cardiac output in female *Gle1*^PFQ/PFQ^ mice but no significant differences in left ventricular systolic function or the structure of the left ventricle (Figure 8B, Supplementary Data File 6-7). Analysis of heart weights also showed an increase in female KI mice, which was absent in males (Supplementary Data File 6-7).

**Figure 8:**
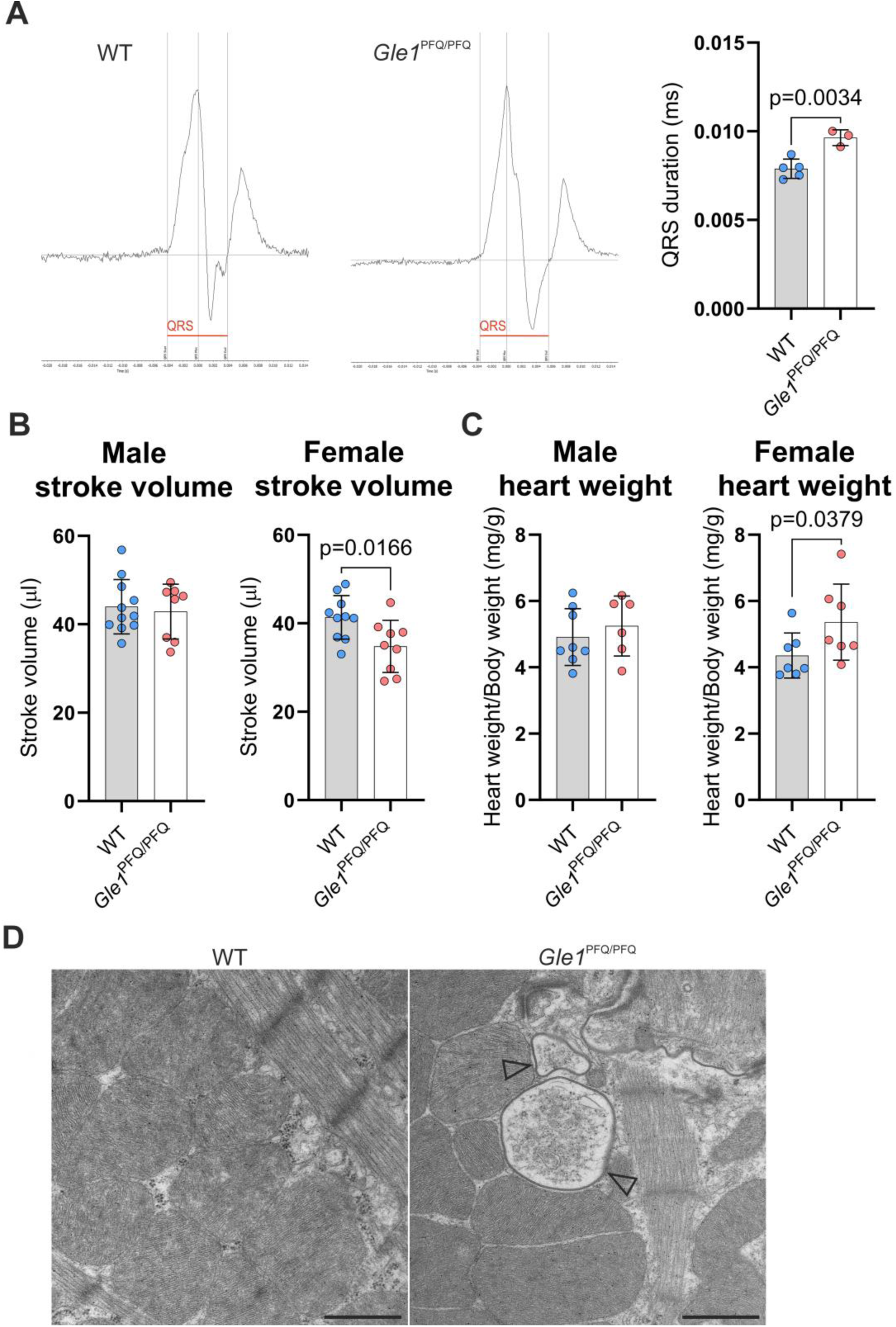
Functional analysis of *Gle1*^PFQ/PFQ^ adult heart. **A)** Representative images of ECG recordings from adult males (8 weeks old). The prolonged QRS wave duration is highlighted by red line. Means ± SD are plotted, as well as individual data points generated from each embryo (WT males: 5, *Gle1*^PFQ/PFQ^ males: 3). Two-tailed Student’s *t* test. **B)** Measurement of left ventricle stroke volume via echocardiography recordings in adult (17-25 weeks old) males and females (WT males: 11, *Gle1*^PFQ/PFQ^ males: 8, WT females: 10, *Gle1*^PFQ/PFQ^ females: 9). Two-tailed Student’s *t* test. **C)** Comparison of heart weight normalized to body weight in adult (17-25 weeks old) males and females (WT males: 8, *Gle1*^PFQ/PFQ^ males: 6, WT females: 7, *Gle1*^PFQ/PFQ^ females: 7). Two-tailed Student’s *t* test. **D)** Representative images from electron microscope analysis of adult mouse left ventricular samples. Arrowheads denote damaged mitochondria (WT females: 3, *Gle1*^PFQ/PFQ^ females: 3). Original magnifications: 6800×. Scale bar: 1 mm.

Characterization of the adult hearts through picrosirius red staining used for myocardial fibrosis detection showed no difference between the genotypes (Supplementary figure 12A), even though the hearts of female KI mice showed mild increase in expression of *Col1a1* and *Ccn2* genes that are associated with fibrosis and tissue remodeling^56,57^ (Supplementary figure 12B). In support of the reduced cardiac output, female KI mice presented an increase in the expression of heart failure related genes^58,59^ *Nppa*, *Nppb*, and *Myh6* while this was not seen in male mice (Supplementary figure 12C). Electron microscopy characterization of female *Gle1*^PFQ/PFQ^ hearts, which exhibited functional and molecular defects, showed occasional mitochondrial defects in otherwise normal cardiac ultrastructure (Figure 8C). Yet, qPCR analysis did not show any irregular expression of tested genes related to mitochondrial modulation^60^: *Pgc-1α1-4*, suggesting no major difference in mitochondrial function or biogenesis between the genotypes (Supplementary figure 12D).

## DISCUSSION

Defects in nucleocytoplasmic transport factors and nucleoporins, including GLE1, are genetic modifiers of disease^61–64^ and specifically *GLE1* mutations are associated with developmental^5^ and neurologic disorders^65^. Previous studies show that most of the nucleoporin knockout mice die during embryogenesis or shortly after birth^66^. We report here that *Gle1* inactivation in mouse mimics previous findings and approves the essential nature of nucleocytoplasmic transport in mammalian cells. Our results demonstrated that total loss of GLE1 function results in embryonic lethality at the pre-implantation period and that this gene is essential for early developmental processes, such as cell proliferation and spatial organization. In essence, *Gle1* KO blastocysts fail to converge the inner cell mass into unified population that would localize in one polarized spot of embryo as seen in controls.

Transcriptional profiling of the *Gle1*^-/-^ embryos identified ion channels and transporters as main biological processes affected by *Gle1* inactivation. Electrical properties of the plasma membrane and alterations in the function as well as distribution of ion channels are known to modulate developing embryos from gamete maturation to neuronal differentiation and function^67^. Our results suggest that early compaction and lineage specification in *Gle1*^-/-^ embryos occur without major challenges but the following processes that prepare embryos for implantation and require extensive exchange of adhesive and antiadhesive factors, fluid transport in response to osmotic gradients, and mitosis-driven paracellular water influx^68^ require normal function of GLE1. Differential expression of ion channels, transporters, and adhesion molecules affects their functions but also modulate membrane potential and intracellular signaling via e.g. calcium^69^ required for survival and thriving.

The severe defects and early lethality of *Gle1*^-/-^ embryos suggest that even in the most severe *GLE1* related developmental disorders, the mutations either modify or disturb the physiological GLE1 functions, rather than induce complete loss of function. The most severe phenotype among the known genetic GLE1-related disorders is associated with an intronic *GLE1* variant causing a rare developmental disorder known as LCCS1^70^. Thus, we genocopied the LCCS1 causative *GLE1*_FinMajor_ variant in mice to produce a *Gle1*^PFQ/PFQ^ KI model that mimics the documented outcome of *GLE1*_FinMajor_ induced alternative splicing^5^. The molecular phenotype of the *Gle1*^PFQ/PFQ^ MEFs indicates that the LCCS1 pathology in mammalian cells may be driven by mechanisms other than simply disrupted nucleocytoplasmic shuttling of poly(A)+ RNA^13^. Our results support the importance of GLE1 function in cellular stress response, as also previously suggested by the studies in HeLa cells^71^. With the emerging trends in stem cell-derived differentiations, using human pluripotent cells with the intronic *GLE1*_FinMajor_ mutation and pushing them towards differentiation into a more diversified cell types shall be the future strategy to verify the current knowledge and uncover the full spectrum of molecular defects resulting in LCCS1 etiology.

*GLE1* variants enriched in isolated populations are associated with CAAHD and LCCS1 disorders^5^. Perinatal death in CAAHD is caused by postnatal respiratory failure, but the cause of fetal death in LCCS1 remains unknown. LCCS1 fetuses show hydrops, pulmonary and skeletal muscle hypoplasia, micrognathia and arthrogryposis, defects in anterior horn motor neurons, and severe atrophy of the ventral spinal cord^5^. Our *Gle1*^PFQ/PFQ^ mouse model presents a milder phenotypic profile and better survivability than what is documented in human patients^72^. It remains to be elucidated whether the mice are less sensitive or more resistant towards the PFQ amino acid insertion due to the interspecies differences in development and physiology^73–75^, or the alternative splicing events induced by the *GLE1*_FinMajor_ mutation are more complex than what has been described by Sanger sequencing of the LCCS1 patient cDNA at the mutation site^5^. With the progress in long-read sequencing methods^76^, checking the full *GLE1* transcript length and amino acid composition in LCCS1 patient cells should be the next step for better understanding of molecular biology in LCCS1 disease and to improve the efforts of modelling this disorder.

Even though the *Gle1*^PFQ/PFQ^ mouse model does not fully phenocopy LCCS1 or other related human diseases, it shows disturbed GLE1 functions at cellular level and thus provides significant insights into the developmental and physiological roles of GLE1 in mammals. Peripheral nervous system defects are reported for LCCS1 patients, who manifest loss of spinal MNs^4,6–8,77^. We show that *Gle1*^PFQ/PFQ^ embryos exhibit a similar, though milder, MN defect than human fetuses and additionally have morphologically shifted MN patterning. Beyond the motoric function, the early MN communication with endothelial cells of growing blood vessels is crucial for spinal cord vascularization and blood vessel patterning^78,79^. Misguided MN patterning can potentially impact the vascular patterning process, affecting the overall integrity of the developing spinal cord. More importantly, we demonstrate that disturbed GLE1 function in the *Gle1*^PFQ/PFQ^ model results in mid-adulthood lethality, irregularly developing paravertebral sympathetic ganglia, and adrenal chromaffin cell abnormalities. Our results additionally show that the embryonic heart innervation is abnormal in *Gle1*^PFQ/PFQ^ mice, which also present mild, gender-specific phenotypic changes in the adult female hearts. Particularly interesting are the transient irregularities in ECG measurements, which could be explained by compensatory or corrective mechanisms during the cardiac maturation process. It is also possible that the *Gle1*^PFQ/PFQ^ mice are more susceptible to stress-induced cardiac arrythmias (not tested here) leading to sudden death of the mice in mid-adulthood.

In conclusion, our findings demonstrate the essential nature of GLE1 functions for early mammalian development and suggest that the pathogenic *GLE1* variants rather modify than completely inactive protein function(s). We provided novel information of modified cell-type specific functions but further studies especially focusing on protein structure are needed for better understanding of how protein interactomes are affected by given pathogenic variant. Additionally, modeling LCCS1 in mouse identified that the importance of GLE1 in developmental biology stretches beyond spinal MNs. The future efforts will reveal if these results can be validated in context of human tissues to enhance understanding of RNA biology in human health and disease.

## Materials and methods

### Mouse generation by CRISPR/Cas9 editing

The FVB/NRj mice were obtained from the Janvier labs (France) and used in the study for genome editing. We confirm that all experiments were performed in accordance with relevant guidelines and regulations. The Committee for animal experiments of the District of Southern Finland approved the animal experiments under the license ESAVI/10982/2021 (valid: 29.4.2021-28.5.2024). The Cas9 protein (HiFi Cas9 nuclease, IDT), gRNA(s), and DNA repair template (Table S1) were microinjected into the zygotes that were further transferred into oviducts of pseudopregnant female. The gRNAs were formed by annealing of the crRNA and universal CRISPR-Cas9 tracrRNA (IDT) and the crRNAs were designed via Benchling (https://benchling.com). The final concentration of reagents for microinjection was 20 ng/µL per Cas9 protein, 25 ng/µL per gRNA, 25 ng/µL per ssDNA.

#### Generation of *Gle1* KO mice

Two gRNAs were employed to cut out approximately 100 base pairs in exon 1 of the *Gle1* gene. Out of the 467 injected zygotes, 5 founders were born and verified by Sanger sequencing. The founder, whose deletion induced reading frameshift and formation of stop codon in exon 2, was chosen for breeding into next generations.

#### Generation of *Gle1* KI mice

The combination of gRNA and a ssDNA repair template were employed to directly knock-in 9 nucleotides (coding the PFQ amino acids) upstream of exon 4 to mimic the outcome of alternative splicing in LCCS1 disease, since the *GLE1*_FinMajor_ is an intronic mutation but mouse and human introns are vastly different. From 54 born pups, 13 founders were identified, 6 of them were verified by Sanger sequencing, and 3 randomly selected founders were bred to the F2 generation. After observing premature lethality of *Gle1*^PFQ/PFQ^ homozygotes amongst progenies of all three founders, only one line was randomly selected for further experiments.

#### Genotyping and DNA sequencing

Mouse pups between 2-3 weeks old were sampled from ear to collect material for genotyping to generate both *Gle1* mouse lines. The DNA was extracted using the Extracta DNA Prep for PCR (95091-025, Quantabio) and used as a template for the PCR reaction, amplified by DreamTaq DNA Polymerase (EP0705, Thermo Fisher) using the SimpliAmp™ Thermal Cycler (A24811, Thermo Fisher). Enzyme digestion of the PCR product by BceAI (R0623L, NEB) had been used for the *Gle1* KI mice to distinguish the carriers of the insertion. Genotype was distinguished from the mixture of DNA fragments on SYBR™ Safe (S33102, Thermo Fisher) stained agarose (BIO-41025, Bioline) gel after electrophoresis. The genotyping primers (supplementary figure 2) have also been used for Sanger sequencing by employing the Eurofins DNA sequencing services (https://www.eurofins.com). The DNA sequences have been viewed and analyzed through the SnapGene software (https://www.snapgene.com).

#### Early embryo genotyping

Either fresh or post-fixed embryos after imaging were washed with PBST and lysed in 10 µL of blastocyst lysis buffer (10 mM Tris-HCl, pH 8.0, 50 mM KCl, 2.5 mM MgCl2, 0.1 mg/ml gelatin, 0.45% Igepal, 0.45% Tween-20, 0.2 mg/ml proteinase K) at +56°C for 30min. Proteinase K was inactivated at 95°C for 10min. Lysate (2-6 µl) was used as a template for PCR for the genotyping.

### Early embryo collection

6-8-week-old heterozygous females were super ovulated by pregnant mare serum - human chorion gonadotrophin (5 IU each) combination^80^ and mated with heterozygous males. Embryos (E2.5, E3.5 or E4.5) were flushed out from the oviducts in M2 medium, and either snap-frozen (RNA isolation), or immediately fixed with 4% PFA for 30 min, or cultured in KSOM medium in a 5% CO_2_ humidified incubator at 37°C: 2-cell embryos until E3.25 and E3.5 for one day (E3.5 + 1) prior to fixation.

### Spinal cord processing for motoneuron analyses

E11.5 embryos were collected after *Gle1*^PFQ/+^ heterozygote-to-heterozygote mating, fixed in 4% PFA in PBS overnight, and genotyped from the tails. The embryos of desired genotype were cryoprotected in 30% sucrose in PBS for 48 hours, before embedding in Tissue-Tek O.C.T. compound on dry ice. The embryos were cryo sectioned using the Leica CM3050 S cryostat at the 16 µm thickness and the brachial region spinal cord sections we collected using the anatomical cues.

### RNA extraction

MEFs were washed with ice cold PBS twice and cells were lysed by TRIzol (15596018, Thermo Fisher) for 5 min on ice. RNA was isolated using the standard phenol-chloroform extraction.

Tissues were quickly dissected, snap frozen in liquid nitrogen, and homogenized in TriZol using the Precellys lysing kit (CK28, Bertin Instruments). The homogenate was left on ice for 5 min, before pushing the content through a 25G needle several times and leaving it on ice for another five minutes. RNA was isolated using the standard phenol-chloroform extraction.

Blastocysts (E3.5) RNA was extracted with Allprep DNA/RNA Micro kit (Qiagen, #80284) according to the instructions. Isolated DNA was used for the genotyping.

### Bulk RNA sequencing of blastocysts

RNA amplification (SMARTer kit), mRNA library preparation, RNA sequencing, and data processing were performed by Novogene (Cambridge, UK) on Illumina Sequencing PE150. Each sample represents a pool of 4 blastocysts (WT, *n*=3; KO, *n*=3). Genes with adjusted p-value < 0.05 and log2(FoldChange) > 1 were considered as differentially expressed.

### Reverse transcription-quantitative polymerase chain reaction (RT-qPCR)

The cDNA was synthesized using the SuperScript IV Reverse Transcriptase (18090050, Thermo Fisher) and diluted in nuclease free water. For the qPCR reaction, 2.5 µl of cDNA were mixed with 1 µl each of forward and reverse primer (Table S2, S3) along with 1.5 µl of PCR water and 5 µl SensiFAST™ SYBR® No-ROX Kit (BIO-98020, Bioline) or with TaqMan probes (Table S4). qPCR was carried out at CFX384 Touch Real-Time PCR Detection System (Bio-Rad). qPCR was performed with at least three biological replicates in two technical replicates. The gene expression was normalized to geometric mean of *Actb*/*Ywhae* or 18s ribosomal RNAand the relative gene expression was calculated using the 2^−ΔΔCT^ method^81^.

### Western blot

Cells were washed with ice cold PBS twice and lysed using 1x RIPA cell lysis solution supplemented with Pierce™ Protease Inhibitor Tablets, EDTA-free (A32965, Thermo Fisher) and Pierce™ Phosphatase Inhibitor Mini Tablets (A32957, Thermo Fisher). The adult heart was snap frozen in liquid nitrogen and homogenized in 1x RIPA with the Precellys lysing kit. The homogenate was pushed through the 25G needle several times, centrifuged at 10 000 RPM for 15 minutes, and the supernatand was collected for analysis. Protein quantity estimation was performed using Pierce™ BCA Protein Assay Kit (23225, Thermo Fisher). In summary, 12.5 µg heart lysate was used for Gle1 quantification, 7.5 µg of MEFs lysate was used for integrated stress response analysis, and 18 µg of protein was employed for nascent protein synthesis quantification in MEFs. Protein was loaded on to in-house prepared 10-15% SDS-PAGE separating gels and 6% stalking gel and separation was performed at 110V at room temperature. Transfer was carried out using the wet blot transfer at 250 mA for 70 minutes onto the nitrocellulose membrane. The membrane was blocked with 5% BSA solution for 2 hours and all the primary antibodies (Table S5) were incubated for overnight at 4°C, while secondary antibodies were incubated for 90 minutes at room temperature. The detection was performed using the WesternBright ECL HRP substrate on Bio-Rad ChemiDoc™ Imaging System. The proteins were either normalized to β-actin after the antibodies were stripped away by the Restore™ PLUS Western Blot Stripping Buffer (46430, Thermo Fisher) for 10 minutes at room temperature and re-blocking and re-staining, or to the total protein load quantified by the Ponceau S (78376, Merck) staining.

### Long range RNA sequencing

RNA isolated from P0 WT and *Gle1*^PFQ/PFQ^ hearts was used for PacBio SMRT sequencing to analyze the presence of any undesired modifications or changes in length of the mouse *Gle1* KI transcript. Sample purity was checked with NanoDrop (ND2000USCAN, Thermo Fisher) and the RNA integrity with Agilent 2100 Bioanalyzer. The library has been prepared from 150 µg of RNA using the SMRTbell prep kit guideline. The quality control, library preparation, and sequencing have been performed by Novogene (UK). Following the sequencing, the demultiplexed files were further processed using PacBio’s IsoSeq bioconda package (v. 4.0.0) following tool documentation provided by Pacific Biosciences. In short, any remaining primer sequences were removed from sample spesific HiFI-reads using lima tool in isoseq mode. Next, ‘isoseq refine’ was used to trim Poly-A tails and to identify and remove concatamers. Isoform-level clustering was done using ‘isoseq cluster2’ function, which was followed by alignment to reference genome using pbmm2, a A minimap2 SMRT wrapper for PacBio data. Reference genome and annotations used were from GENCODE release M33. ‘Isoseq collapse’ was used to collapse redundant transcripts into unique isoforms using recommended settings for bulk tissue IsoSeq. Pigeon PacBio Transcript Toolkit was then used to prepare the input files and then to classify transcripts against given annotations.

### Tissue collection and processing

#### Paraffin sections

Tissues were collected at the stages indicated in the text. Samples were fixed overnight in 4% paraformaldehyde (PFA), processed and embedded in paraffin using an automated tissue processor Tissue-Tek (Sakura), cut into 5-μm-thick sections, and allowed to dry overnight. Immunofluorescent staining of paraffin sections was performed following standard protocol with xylene-alcohol deparaffination series followed by heat-induced antigen retrieval in 20 mM Tris-HCl, 1 mM EDTA buffer at pH 8.5. After cooling to room temperature, tissue sections were blocked 1 h in 50 mM Tris-HCl, 100 mM NaCl, 0.1% Tween-20, at pH 7.5 (TNT) supplemented with 10% FBS, followed by incubation with primary antibodies at 4°C overnight, TNT washing, and 1 h incubation with secondary antibodies. Additionally, fluorescent dye Hoechst 33342 (H3570, Thermo Fisher) was used at 1:1000 dilution for counterstain.

#### Frozen sections

Brachial spinal cord frozen sections were thawed for 5 minutes at +4°C and for 10 minutes at the room temperature, followed by washing and rehydration using PBS for 2×10 minutes. Sections were permeabilized for 30 minutes using 0.5% Triton X-100 in PBS and blocked for 1 hour with 5% FBS in permeabilization buffer. Subsequently, sections were incubated with primary antibodies at 4°C overnight in blocking buffer, washed in PBS 3x, and 1 h incubation with secondary antibodies in blocking buffer. Additionally, fluorescent dye Hoechst 33342 (H3570, Thermo Fisher) was used at 1:1000 dilution for counterstain.

#### E14.5 limb innervation

The whole forelimb was cut off with scissors, fixed overnight in 4% PFA, permeabilized for 30 minutes using 0.5% Triton X-100 in PBS, and blocked for 1 hour with 5% FBS in permeabilization buffer. The limbs were incubated in primary antibody, diluted in the blocking buffer, for 48 hours at 4°C. Afterwards, limbs we washed in 0.1% Triton X-100 in PBS for 3×15 minutes, and incubated in secondary antibody diluted in the blocking buffer overnight at 4°C. Following up with washing the limbs again, samples were dehydrated with ethanol series (pH=9, at 4°C, gently shaking): 30% for 2 hours, 50%, 70%, 2× 100% for 4 hours each, finished with 100% ethanol overnight. After dehydration, the samples were transferred to ethyl cinnamate (ECi, 112372; Merck) for at least 6 hours before imaging, using the ECi also as a mounting medium.

### Immunostaining

#### Early embryos

Fixed embryos were permeabilized with 0.3% Triton X-100/0.1M glycine in PBS for 20min at room temperature and washed with PBS + 0.1% Tween-20 (PBST). The embryos were blocked with 10% filtered FBS in PBST for 30min and incubated with primary antibodies diluted in 10% FBS in PBST + 0.2% NaN_3_ at +4 °C overnight. After two washes with PBST, embryos were incubated with secondary antibodies for 2h at room temperature. After a washing step, nuclei were stained with Hoechst for 8min. Individual embryos were mounted in an imaging chamber in 30% glycerol in PBS, and immediately imaged. List of antibodies is summarized in Table S6.

#### MEFs

20 000 cells were seeded and grown on fibronectin coated coverslips until desired confluence. Cells were fixed with 4% PFA in PBS for 20 min at room temperature and washed twice with wash buffer (0.2% BSA in PBS + 0.1% Tween 20). The cells were permeabilized with 0.1% Triton X-100 + 50mM Glycine for 15 minutes, washed with wash buffer three times while on horizontal shaker, blocked with 5% FBS in 0.1% Triton X-100 in PBS to block unspecific binding and incubated with primary antibodies (Table S6)diluted in the blocking buffer overnight at 4°C. Cells were washed with the wash buffer three times again before incubation with the secondary antibodies diluted in the blocking buffer for 90 min. Following another washing step, the nuclei were stained with Hoechst for 8min. After washing the cells with PBS and mounted on glass slides using the Epredia™ Immu-Mount™ (Thermo Fisher). Optionally, phalloidin staining has been added after the secondary antibody staining by diluting the phalloidin in PBS and staining for 2 hours.

#### Heart innervation

E16.5 heart ventricles were dissected, fixed with 4% PFA in PBS overnight and then immunostained and clarified using a published protocol^82^. First, the hearts were incubated in CUBIC reagent-1 solution containing 25 % (wt/vol) of urea, 25 % (wt/vol) of N,N,N,N-tetrakis(2-hydroxypropyl)ethylenediamine and 15 % (wt/vol) Triton X-100 in water. Tissues were immersed in the solution (600 ul in a 4-well plate) and gently rocked at 37°C for 5 hours. Hearts were blocked with 5% FBS in PBS, 0.1% triton X-100 for 12 hours at room temperature and incubated with primary antibody in the blocking buffer overnight at +4°C. The tissues were washed with 0.5% Tween 20 in PBS for 3×10 min before incubating the tissues in secondary antibody in blocking buffer. After washing the hearts for 3×10 min again, they were equilibrated in 20% sucrose in PBS overnight. Subsequently, hearts were incubated in Cubic reagent-2 (50% sucrose, 25 % (wt/vol) of urea, 10 % (wt/vol) of triethanolamine, 0,1% triton X-100) for 1 day in dark and at room temperature. Cubic reagent-2 was also employed as a mounting medium for imaging.

#### Paravertebral sympathetic ganglia

E14.5 embryos were collected and cut beneath the thoracic cavity in half before removing the liver. The upper half of the embryo was fixed with 4% PFA in PBS overnight and then immunostained and clarified using a slightly modified iDISCO protocol^83^. Fixed samples were washed in PBS for 1 hour twice, then in 50% methanol (in PBS) for 1 hour, 80% methanol for 1 hour, 100% methanol for 1 hour twice. Samples were then bleached with 5% H_2_O_2_ in 20% DMSO/methanol (1 vol 30% H_2_O_2_/1 vol DMSO/4 vol methanol, ice cold) at 4°C overnight. After bleaching, samples were washed in methanol for 1 hour twice, then in 20% DMSO/methanol for 1 hour twice, then in 80% methanol for 1 hour, 50% methanol for 1 hour, PBS for 1 hour twice, and finally in PBS/0.2% Triton X-100 for 1 hour twice before further staining procedures. Pretreated samples were incubated in PBS/0.2% Triton X-100/20% DMSO/0.3 M glycine at 37°C overnight, then blocked in PBS/0.2% Triton X-100/10% DMSO/6% FBS at 37°C overnight. Samples were washed in PBS/0.2% Tween-20 with 10 μg/ml heparin (PTwH) for 1 hour twice, then incubated in primary antibody dilutions in PTwH/5% DMSO/3% FBS at 37°C for 5 days. Samples were then washed in PTwH for 1 day, then incubated in secondary antibody dilutions in PTwH/3% DMSO/3% FBS at 37°C for 4 days. Samples were finally washed in PTwH for 2 days before clearing and imaging. Immunolabeled tissues were incubated overnight in 10 ml of 50% v/v tetrahydrofuran/H2O (THF) (186562-12X100ML, Merck). Samples were then incubated for 1 hour in 10 ml of 80% THF/H2O and twice 1 hour in 100% THF and then in dichloromethane (DCM) (270997-12X100ML, Merck) until they sank at the bottom of the vial. Finally, samples were incubated in 15 ml of dibenzyl ether (DBE) (108014-1KG, Merck) until clear (∼2 hours) and then stored in DBE at room temperature. Before imaging, the sample were incubated in ethyl cinnamate for 4 hours at room temperature (112372-100G, Merck).

### Mouse embryonic fibroblast isolation, growth, and functional analyses

Plugged females from *Gle1*^PFQ/+^ heterozygote to heterozygote mating were euthanized by CO_2_ at E13.5 stage. Uterus was removed and the embryos dissected in Petri dishes. The head was cut off and the organs removed before washing the embryos in PBS to remove blood as much as possible. The carcasses were minced in PBS into cubes of about 2-3 mm in diameter with scissors. The cubes were incubated in trypsin/EDTA (25200-056, Thermo Fisher) at 37°C for 30 minutes with stirring, with additional DNase I (10649890, Thermo Fisher) and trypsin/EDTA added for further 30-minute incubation. Cell suspensions were centrifuged at 1000 rpm for 5 minutes. The cell pellets were washed with MEFs culture medium (DMEM (31966-021, Gibco) + 10% FBS (10500064, Thermo Fisher) + penicillin/streptomycin (15140122, Thermo Fisher)) twice to eliminate the trypsin activity and finally resuspended in the culture medium. Living nucleated cells were seeded in 30 mm tissue culture dishes and expanded until passage 2 (P2), before freezing the trypsinized cells in the freezing media (7.5% DMSO and 92.5% fetal bovine serum) until later use. The preserved MEFs with desired genotype were thawed before experiment and expanded. All of the experiments with MEFs were performed at passage 4 (P4) and at 60-70% confluence on glass fibronectin (1030-FN-05M, R&D Systems) coated coverslips or plastic culture plates (Starlab).

#### poly(A)+ RNA distribution

Fluorescent *in situ* hybridization of the poly(A)+ RNA has been performed on MEFs grown on fibronectin coated coverslips. Cells were fixed with 4% PFA in PBS for 20 min at room temperature, washed in PBS twice, permeabilized with ice cold methanol for 10 min and then incubated in 70% ethanol at 4°C overnight. On the next day, the cells were incubated for 5 min at room temperature in wash buffer (10% formamide in 2× saline sodium citrate (SSC)). Then, the coverslips were transferred face down onto a drop of 100 μl of hybridization buffer (10% formamide (AM9342, Thermo Fisher), 2× SSC, 1 mg/ml yest tRNA (AM7119, Thermo Fisher) and 10% Dextran sulfate (J14489, Thermo Fisher) containing 100 nM probe (5’-Cy3-oligo-dT(30), IDT), in a humidified chamber sealed with parafilm and incubated in the dark at 37°C overnight. After hybridization, the coverslips were transferred face up to a fresh well and washed twice in wash buffer in the dark at 37°C, for 30 min each wash, and then washed briefly in 1× PBS. Hoechst (10 min) was used as a DNA counterstain.

#### RNA synthesis

Global RNA synthesis was analyzed in MEFs grown on fibronectin coated coverslips through metabolic labeling. Cells were treated with 1 mM ethynyl uridin (EU) for 1 hour before fixation with 4% PFA in PBS for 20 min. Cells were washed and permeabilized with 0.5% Triton X-100 in PBS and click chemistry was performed according to the Click-iT™ RNA Alexa Fluor™ 594 Imaging Kit instructions.

#### Protein synthesis

*De novo* protein synthesis in MEFs grown on 10 cm plates was analyzed through the analysis of puromycilated proteins. The cell culture medium was replenished 2 hours before the experiment. Cells were treated with 10 μM puromycin (A1113803, Thermo Fisher) for 30 min. After puromycin pulse, cells were washed with ice-cold PBS containing 100 μg/mL cycloheximide (C-0934, Merck) to inhibit further puromycin labeling. Samples were analysed by western-blot and puromycin incorporation was visualized by using the mouse monoclonal antibody MABE341 (1:25 000) for immunolabeling and chemiluminiscent detection.

### Cell cycle and apoptosis analyses

#### Flow cytometry

MEFs were grown in 10cm plates until 60-70% confluence before washing with PBS and trypsinization (15090046, Thermo Fisher). Cells were fixed by slowly adding ice-cold 70% ethanol and keep at 4°C for 4 hours. Cells were centrifuged (500 RPM for 5 min) and the pellet was washed twice with 2% FBS in PBS. Cells were treated with 100 µL of RNase A for 1×10^6^ cells at a concentration of 1 mg/mL to the cells and incubated at 37°C for 30 min. MEFs were stained with propidium iodide (421301, BioLegend). Samples were analyzed BD Accuri C6 plus Analyser, cell population gated in the scatter plot and the cell cycle viewed as FL3-H using a linear scale.

#### Proliferation assay

EdU labelling was performed using Click-iT EdU Alexa Fluor 488 Imaging Kit (C10337, Thermo Fisher) according to the manufacturer’s instructions. Blastocysts (E3.5) were incubated with 10 µM EdU in KSOM media in a 37°C incubator (5% CO_2_) for 30 min while MEFs were incubated for 6 hours in the standard growth medium, before fixation with 4% PFA in PBS for 30 min.

#### TUNEL assay

DNA damage in the blastocysts (E3.5) was assessed by using the In Situ Cell Death Detection Kit, Fluorescein (11684795910, Merck) according to the manufacturer’s instructions.

### SA-β-Gal staining

MEFs were seeded at passage 4 on the 6-well plates and grown for 48 hours until 60-70% confluence. The SA-β-Gal staining was performed using the Senescence Detection Kit (ab65351, Abcam). Cells were washed with PBS once, fixed with the kit fixative for 15 min, and stained with X-gal (10 µg/mL) for 12 hours at 37°C (incubator without CO_2_).

### Adult heart characterization

#### ECG and HRV measurements

The anesthetized mouse was placed on a platform with its paws attached to three paw sized electrodes. Gel was applied to improve conductivity. One channel ECG-signal was measured from right paw to left paw (lead II). The ECG-system was connected to the laptop with an analog to digital transformer with 10kHz sampling frequency (Power lab/8SP, ADInstruments, Australia) managed with Labchart software v7.3.2, ADInstruments, Australia). Ectopic and abnormally shaped beats were removed from the analysis. ECG samples were analyzed manually by a trained operator, who identified the P-wave onset and offset, QRS boundaries and peak amplitudes of P, R and S waves. For the HRV analysis, ECG was recorded for a 5-min period. HRV parameters were analyzed and calculated by the Labchart software tool “HRV analysis”.

#### Echocardiography

High resolution ultrasound system Vevo 2100 (Fujifilm-Visualsonics, Toronto, Canada) was used for the image acquisition with MD-550 linear array transducer (40MHz, 30 um axial, 90 um lateral resolution). Transthoracic echocardiography was performed on unconscious, anesthetized mice (isoflurane inhalation, 4% for induction and 1.7% for maintenance) on a warm examination platform. Morphological data (interventricular septum thickness (IVS), left ventricular internal diameter (LVID) and left ventricular posterior wall thickness (LVPW)) were collected and parameters for systolic function were calculated with the formulas: fractional shortening = ((LVID;d-LVID;s)/LVID;d)x100, ejection fraction = ((LV Vol;d-LV Vol;s)/LV Vol;d)x100, LV Vol = (7.0 / (2.4+LVID)) x LVID3. The image analysis was performed with Vevo Lab 5.8.1 software by an experienced sonographer blinded to the genotype.

#### Analysis of myocardial fibrosis

To quantify myocardial collagen content, the sections were stained with Picro-Sirius Red (Picro-Sirius Red Solution ab246832, Abcam) as described previously^84^. The sections were scanned with a brightfield digital slide scanner (40x objective, NanoZoomer S60, Hamamatsu) and the obtained images were saved to an external hard drive. Collagenous tissue was quantified automatically with a specially tailored script within Visiopharm software utilizing bayesian quadratic discriminant analysis of red-green contrast to detect and calculate the area of collagenous and non-collagenous tissue.

#### Transmission electron microscopy

Heart samples were fixed in 1% glutaraldehyde and 4% formaldehyde in 0.1 M phosphate buffer, postfixed in 1% osmium tetroxide, dehydrated in acetone and embedded in Epon LX 112 (Ladd Research Industries). Thin sections were cut with a Leica UC6 ultramicrotome, stained in uranyl acetate and lead citrate and examined by Tecnai G2 Spirit 120 kV transmission electron microscope. The images were captured using a Quemesa CCD camera (EMSIS GmbH, Muenster, Germany).

### Image acquisition

Early embryos were imaged with Leica TCS SP8 X (z-stack: 1µm per step) confocal equipped with the Leica HyD hybrid detector and HC PL APO CS2 63x/1.40 objective at 512×512 or 1024×1024 format. For fluorescence intensity analysis, MEFs were imaged with the Zeiss Axio Imager (z-stack: 7 sections, 1 µm per step) equipped with Hamamatsu Orca Flash 4.0 LT B&W camera and using the EC Plan Neofluar 40x/1.3 and 63x/1.4 objectives. For stress granule analysis, the stressed out MEFs were imaged via Leica TCS SP8 X (z-stack: 10 sections, 0.5 µm per step) confocal equipped with the Leica HyD hybrid detector and HC PL APO CS2 63x/1.40 objective at 3144×3144 format. Images were then deconvoluted using the HyVolution pipeline. E16.5 heart ventricle whole-mounts were imaged by the Andor Dragonfly spinning disc microscope (z-stack: 900 sections, 1 µm per step + tile scan: 2×2) with Zyla 4.2 sCMOS camera and Plan Fluor DL 10x/0.3 objective. The images were stitched and deconvoluted in the Leica LAS X 4.6.0 software. The paravertebral sympathetic ganglia analysis was performed using the LaVision Biotec Ultramicroscope II Light Sheet microscope with the Andor Neo 5.5-CL3 camera (zoom: 0.8, thickness: 6.69 µm, sheet NA: 0.035, step size: 3.35 µm). Following the SA-β-Gal staining, representative images of SA-β-Gal-positive MEFs were acquired with the Leica DMi1 at the 10x magnification. Furthermore, Hoechst-stained nuclei were imaged and overlayed with SA-β-Gal stain visible in the brightfield using the Leica TCS SP8 X with the HC PL APO CS2 10x/0.40 objective. The heart paraffin sections were imaged either with Andor Dragonfly Zyla 4.2 sCMOS camera and Plan Fluor DL 10x/0.3 objective (E16.5 sections) or Zeiss Axio Imager at 40x magnification (adult sections). The E12.5 spinal cords were imaged with Andor Dragonfly Plan Apo λ 20x/0.75 objective (z-stack: 1 µm per step, 2×2 tile scan). The E14.5 forelimbs were imaged with Andor Dragonfly Plan Fluor DL 10x/0.3 objective (z-stack: 1 µm per step, 2×2 tile scan).

### Image analysis

MEFs have been segmented by CellProfiler 4.2.1 either to measure the mean fluorescence intensity of target structures in subcellular compartments or to quantify the number of stress granules as speckle-type objects per cell (threshold at least 4 pixels per speckle). Heart innervation was analyzed using the Imaris 10.0 filament tracer plug in and normalizing the innervated surface area to the total ventricle surface. Paravertebral sympathetic ganglia and early mouse embryos were segmented and analyzed in Imaris as well. SA-β-Gal-positive MEFs were counted manually, after the segmentation of Hoechst positive nuclei, via cell counter plug in in Fiji. The analysis of heart sections through segmentation and thresholding was performed in Fiji. The colocalization of TH and IB4 was performed via JaCoP^85^ plug in in Fiji.

The E12.5 motoneuron soma coordinates were acquired with respect to the midline using Fiji software. To assign x and y coordinates, the maximal height (from the ventral lower border of the spinal cord under the central canal to the dorsal-most border of the spinal cord) and width (from the center of the central canal to the most lateral edge) of each hemi-section were measured for normalization^86,87^. Contour, and density plots were created using the ‘ggplot2’ package in R-4.4.1.

### Statistics and reproducibility

Statistical analyses were performed using GraphPad Prism software (GraphPad, version 9.2.0) or R-4.4.1. Statistical significance was determined by the specific tests indicated in the corresponding figure legends. Only two-tailed tests were used. All experiments presented in the Article were repeated in three independent biological replicates at least. No statistical methods were used to predetermine sample sizes. Data collection and analysis were not performed blinded.

## Data availability

The KO blastocyst RNA-seq data (PRJNA1162070), as well the KI mouse long range RNA sequencing results (PRJNA1162399) have been deposited in NCBI’s Sequence Read Archive (SRA).

## Acknowledgements

The formation of the new *Gle1* mouse lines, MEF isolation, and early embryo flushing were carried out with the support of HiLIFE GM-Unit Core Facility, University of Helsinki, Finland; a member of Biocenter Finland. EM images were made with the help of Biocenter Oulu, Electron Microscopy Core Facility. We also thank the Biomedicum Imaging Unit for microscopy services, and the University of Helsinki Flow Cytometry Unit for technical assistance. This work was supported by Academy of Finland (348906), Jane and Aatos Erkko Foundation (FinnDisMice-project) and University of Helsinki funds.

