## Supplementary for "GLE1 dysfunction compromises cellular homeostasis, spatial organization, and peripheral axon branching"

**Supplementary figure 1**

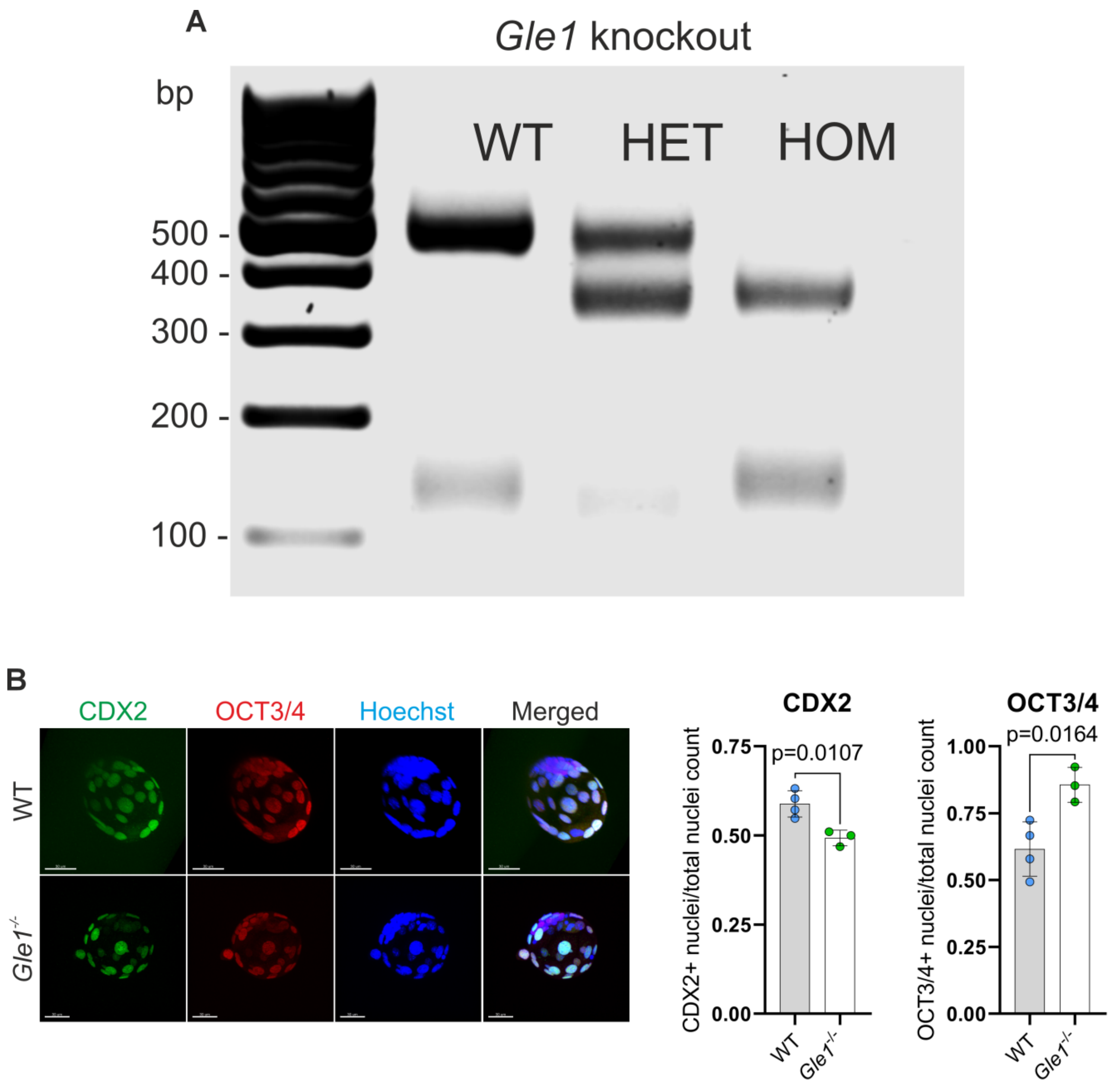


**Supplementary figure 1**: *Gle1* knockout (*Gle1*$^{-/-}$) blastocyst characterization

**A)** Comparison of genotyping results via migration of the PCR amplicons, demonstrating the deletion of the *Gle1* gene.

**B)** Maximum projections of E3.5 embryos flushed from superovulated females, immunostained markers for trophectoderm (CDX2) and inner cell mass marker (OCT3/4) in both WT and *Gle1*$^{-/-}$ early blastocysts. The quantification involves determining the number of CDX2-positive and OCT3/4-positive nuclei relative to the total number of nuclei. Means ± SD are plotted, as well as individual data points generated from each embryo (WT: n=4, *Gle1*$^{-/-}$: n=3). Scale bar 30 µm, two-tailed Student's t test.

**Supplementary figure 2**

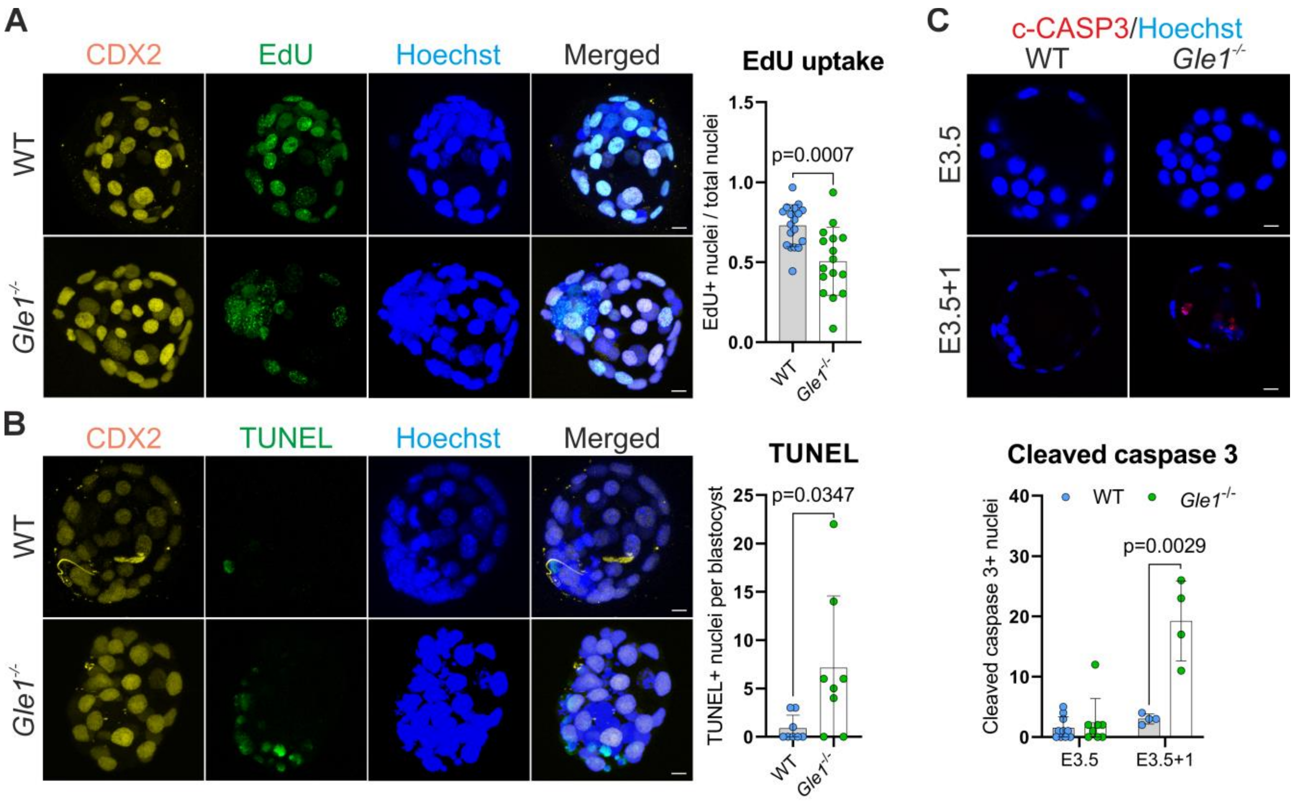


**Supplementary figure 2**: Disrupted cell cycle and cell death in *Gle1* knockout (*Gle1*$^{-/-}$) blastocysts

**A)** Proliferation assay of E3.5 blastocysts and quantification of EdU uptake (10 µM EdU, 30 min) as the number of EdU-positive nuclei relative to the total number of nuclei. Means ± SD are plotted, as well as individual data points generated from each embryo (WT: n=18, Gle1$^{-/-}$: n=16). Scale bar 10 µm, two-tailed Student's *t* test.

**B)** Terminal deoxynucleotidyl transferase dUTP nick end labeling (TUNEL) assay for detecting DNA fragmentation in E3.5 blastocysts. Quantification of DNA damage as the number of TUNEL-positive nuclei per blastocyst. Means ± SD are plotted, as well as individual data points generated from each embryo (WT: n=8, Gle1-/-: n=8). Scale bar 10 µm, two-tailed Student's t test.

**C)** Cross-section of E3.5 and E3.5+1 cultured blastocysts immunostained for cleaved caspase 3 as a marker of apoptosis. Quantification of apoptotic cells as the number of cleaved caspase 3-positive nuclei per blastocyst. Means ± SD are plotted, as well as individual data points generated from each embryo (E3.5 WT: n=10, *Gle1*$^{-/-}$: n=8; E3.5+1 WT: n=4, *Gle1*$^{-/-}$: n=4). Scale bar 10 µm, two-tailed Student's t test.

**Supplementary figure 3**

A

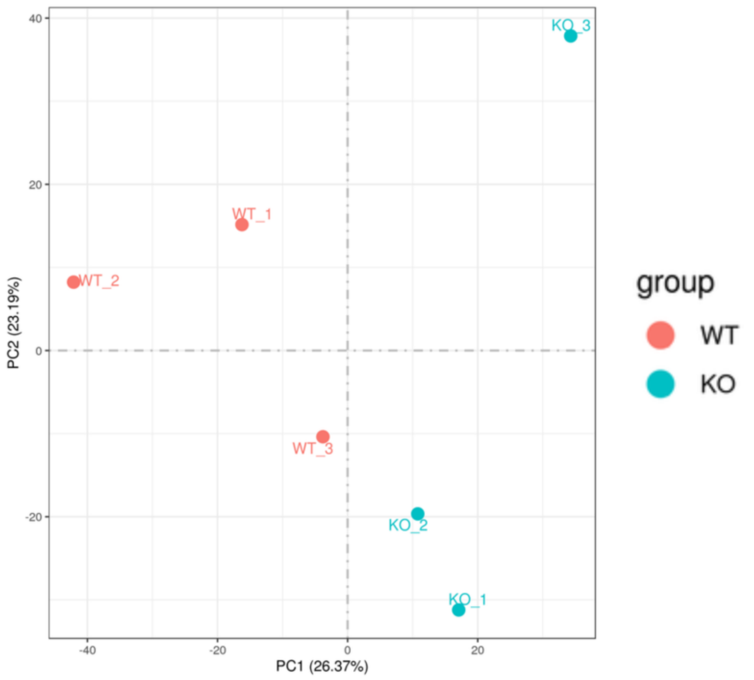


B

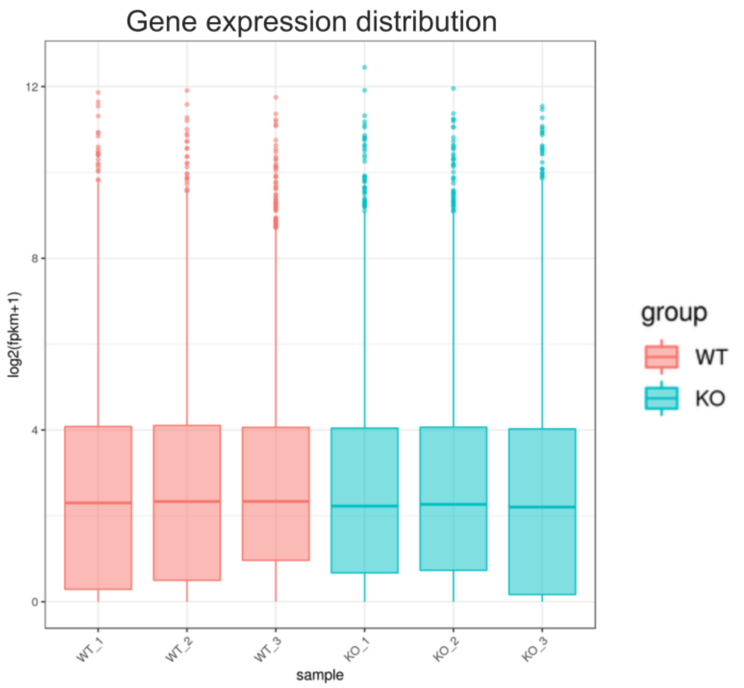


**Supplementary figure 3**: E3.5 blastocyst RNA seq analysis

**A)** Principal component analysis of the whole transcriptome. Points represent the whole transcriptomes of WT (red), and *Gle1*$^{-/-}$ (KO) (blue) E3.5 blastocysts. PC1 represents principal component one, and PC2 represents principal component two, both of which explain the first and the second higher variance among embryos.

**B)** Gene expression distribution boxplot of each sample.

**Supplementary figure 4**

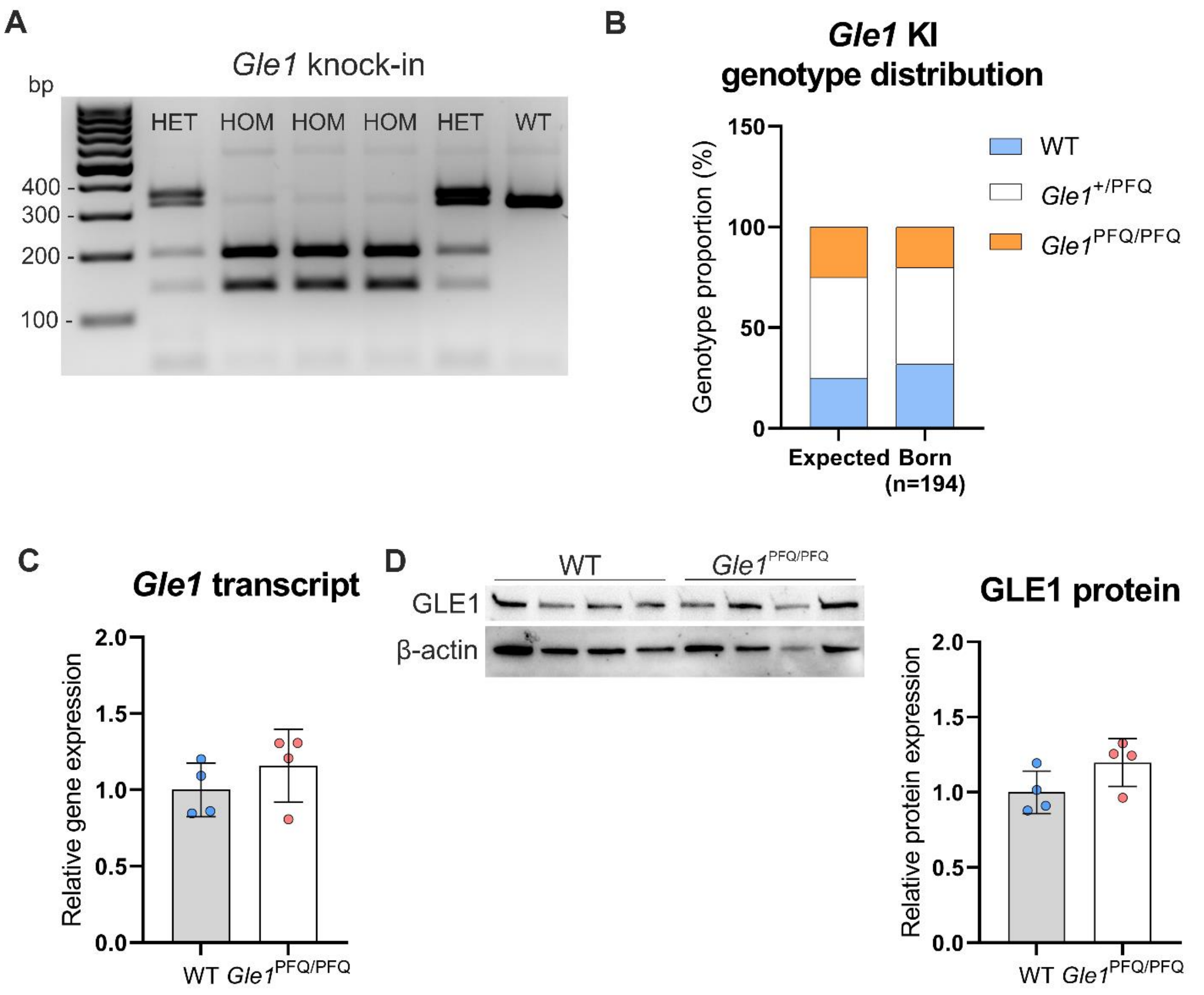


**Supplementary figure 4**: Characterization of GLE1 transcript and protein in *Gle1*$^{PFQ/PFQ}$ mice

**A)** Genotyping results of *Gle1* KI mice by PCR followed by amplicon digestion and analysis of the products on agarose gel.

**B)** Genotype frequencies of mice produced from *Gle1*$^{+/PFQ}$ heterozygote to heterozygote mating (21 litters).

**C)** RT-qPCR quantification of *Gle1* gene expression in 10-week old adult hearts. Means ± SD are plotted, as well as individual data points generated from each mouse (n=4 per genotype), two-tailed Student's *t* test.

**D)** Western blot analysis of GLE1 protein expression in adult heart (10 weeks), normalized to the β-actin expression. Means ± SD are plotted, as well as individual data points generated from each mouse (n=4 per genotype), two-tailed Student's *t* test.

**Supplementary figure 5**

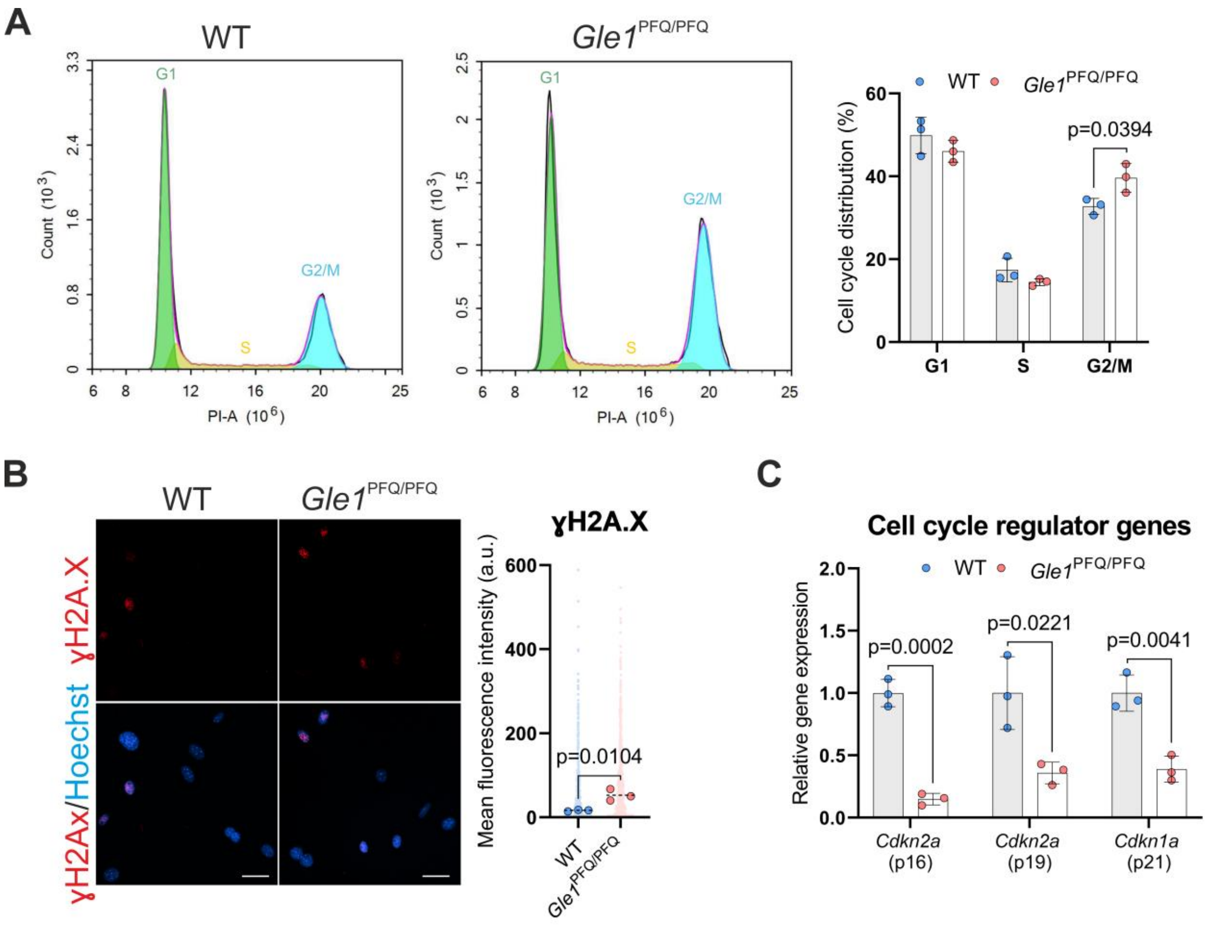


**Supplementary figure 5:** Cell properties of the *Gle1*PFQ/PFQ mouse embryonic fibroblasts (MEFs)

**A)** Distribution of the MEFs in the G1, S and G2/M phases of the cell cycle after 48 h by propidium iodide (PI) staining and flow cytometry. Means ± SD are plotted, as well as individual data points generated from each embryo (three embryos per genotype), two-tailed Student's *t* test.

**B)** Immunostaining and quantification of ɣH2Ax as a marker of DNA damage. Individual data points generated from each cell are plotted (150 cells per embryo, three embryos per genotype), as well as median values from each embryo (large dots). Scale bar 50 µm, two-tailed Student's *t* test.

**C**) RT-qPCR quantification of *Cdkn2a* (p16 and p19) and *Cdkn1a* (p21) mRNA in MEFs at passage 4. Means ± SD are plotted, as well as individual data points generated from each embryo (three embryos per genotype). Two-tailed Student's *t* test.

### Supplementary figure 6

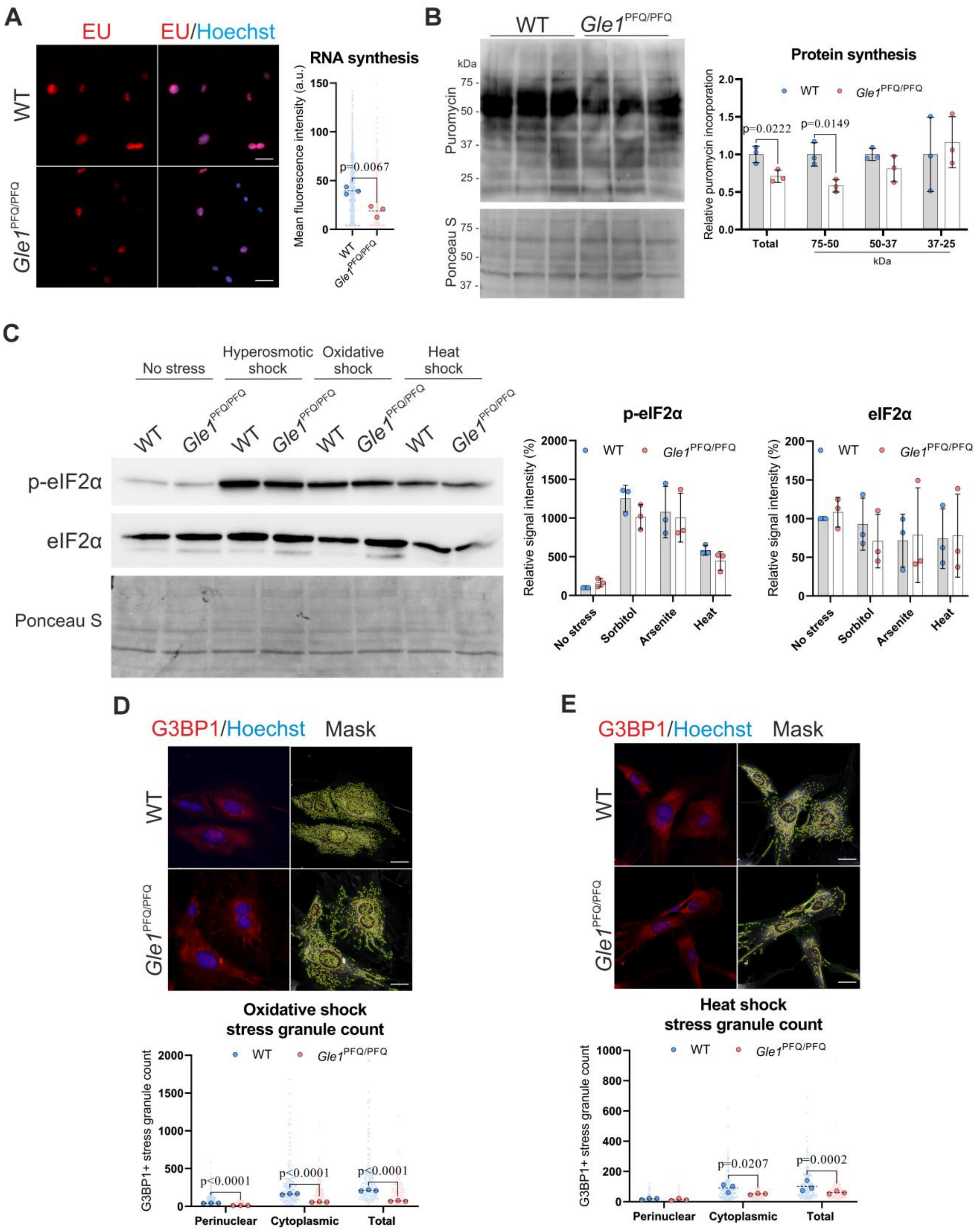


**Supplementary figure 6**: Functional analysis in *Gle1*$^{PFQ/PFQ}$ mouse embryonic fibroblast (MEFs)

**A)** Quantification of nascent RNA synthesis via ethynyl uridine metabolic incorporation (1mM EU, 1 hour pulse). Individual data points generated from each cell are plotted (150 cells per embryo, three embryos per genotype), as well as median values from each embryo (large dots). Scale bar 50 µm, two-tailed Student's *t* test.

**B)** The bulk protein translation was measured by puromycin incorporation (10 µM puromycin, 30-minute pulse) and western blot analysis of the cell lysate, normalized to the total protein load (Ponceau S-stained membrane). Means ± SD are plotted, as well as individual data points generated from each embryo (three embryos per genotype). Two-tailed Student's *t* test.

**C)** The integrated stress response was analyzed by comparing the levels of eIF2α and it phosphorylated form p-eIF2α between wild type and knock-in MEFs upon different stress conditions: no stress vs 60 minutes of hyperosmotic shock (sorbitol)/oxidative shock (sodium arsenate)/heat shock (43 °C), and normalization of the total protein load (Ponceau S-stained membrane). Means ± SD are plotted, as well as individual data points generated from each embryo (three embryos per genotype).

**D)** Maximum projection of MEFs after stress challenge with oxidative shock (500µM sodium arsenate, 1 hour) and stress granule quantification as G3BP11 positive speckle type objects (yellow) in the cytoplasm (green) and around the nucleus (red). Individual data points generated from each cell are plotted (50 cells per embryo, three embryos per genotype), as well as median values from each embryo (large dots). Scale bar 30 µm, two-tailed Student's *t* test.

**E)** Maximum projection of MEFs after stress challenge with heat shock (43°C, 1 hour) and stress granule quantification as G3BP1 positive speckle type objects (yellow) in the cytoplasm (green) and around the nucleus (red). Individual data points generated from each cell are plotted (50 cells per embryo, three embryos per genotype), as well as median values from each embryo (large dots). Scale bar 30 µm, two-tailed Student's *t* test.

**Supplementary figure 7**

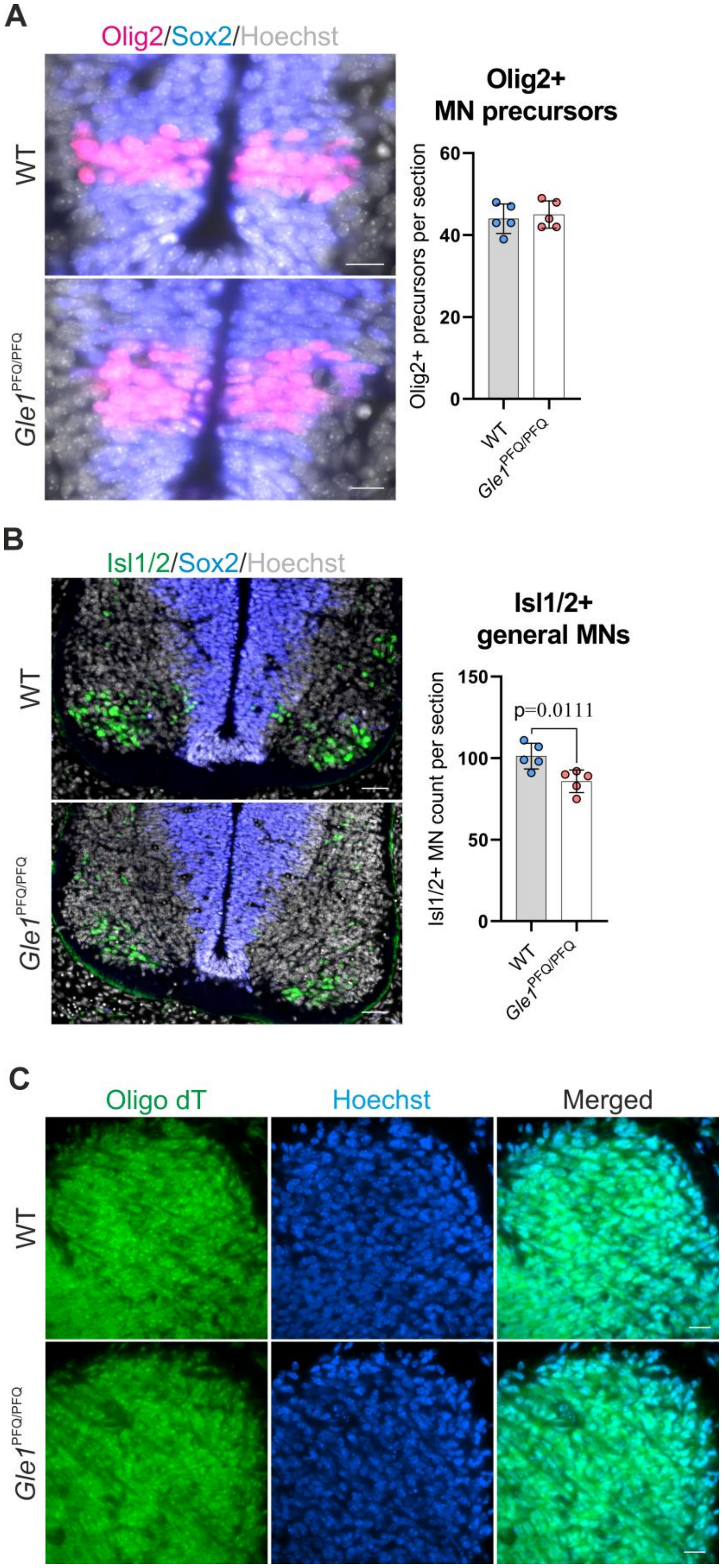

**Supplementary figure 7**: Motor neuron (MN) development in E11.5 *Gle1*$^{PFQ/PFQ}$ mice

**A)** Transverse sections through the brachial level of *Gle1*$^{PFQ/PFQ}$ embryos quantified for the immunostained OLIG2-positive progenitors (pink) of MNs from amongst the SOX2-positive neuroepithelial cells (blue) of the neural tube, counterstained with Hoechst (grey). Individual data points generated from each mouse embryo are plotted (average value from 2 analyzed sections per embryo, five embryos per genotype). Scale bar 20 µm, two-tailed Student's *t* test.

**B)** Transverse sections through the brachial level of *Gle1*$^{PFQ/PFQ}$ embryos quantified for the immunostained Isl1/2-positive general MNs (green) at the lateral motor column, counterstained with SOX2 (blue) and Hoechst (grey). Scale bar = 50µm. Individual data points generated from each mouse embryo are plotted (average value from 2 analyzed sections per embryo, five embryos per genotype). Scale bar 20 µm, two-tailed Student's *t* test.

**C)** Representative images of oligo dT (green) hybridized lateral motor columns in wild type (WT) and *Gle1*$^{PFQ/PFQ}$ mice, demonstrating normal poly(A)+ RNA distribution between genotypes in the embryonic spinal cords (five embryos per genotype). Scale bar 20 µm

**Supplementary figure 8**

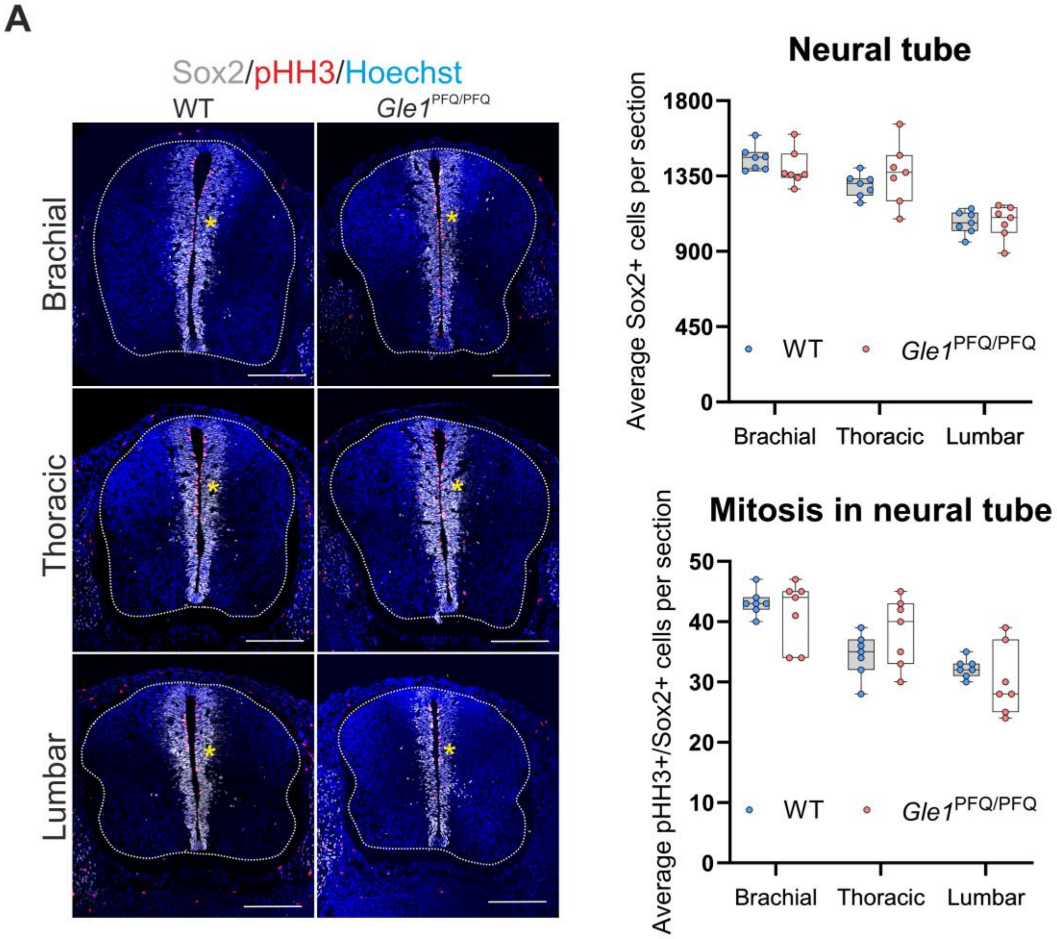


**Supplementary figure 8**: Quantitative analysis of the developing spinal cord in E12.5 *Gle1*$^{PFQ/PFQ}$ embryos

**A)** Representative images of E12.5 spinal cord transverse sections at the brachial, thoracic, and lumbar regions, immunolabelled for SOX2 (white) as a marker of neural progenitors constituting the ventricular zone (yellow asterisk) of the developing neural tube and pHH3 (red) as a marker of the mitotic cells. White borderline demarcates the spinal cord shape. Box and whisker plots, as well as individual data points (n=7) generated from each mouse (i.e. the average edge length/section area/cell count from multiple sections – brachial: 3, thoracic: 4, lumbar: 4), two-tailed Student's t test Scale bar 200 µm.

**Supplementary figure 9**

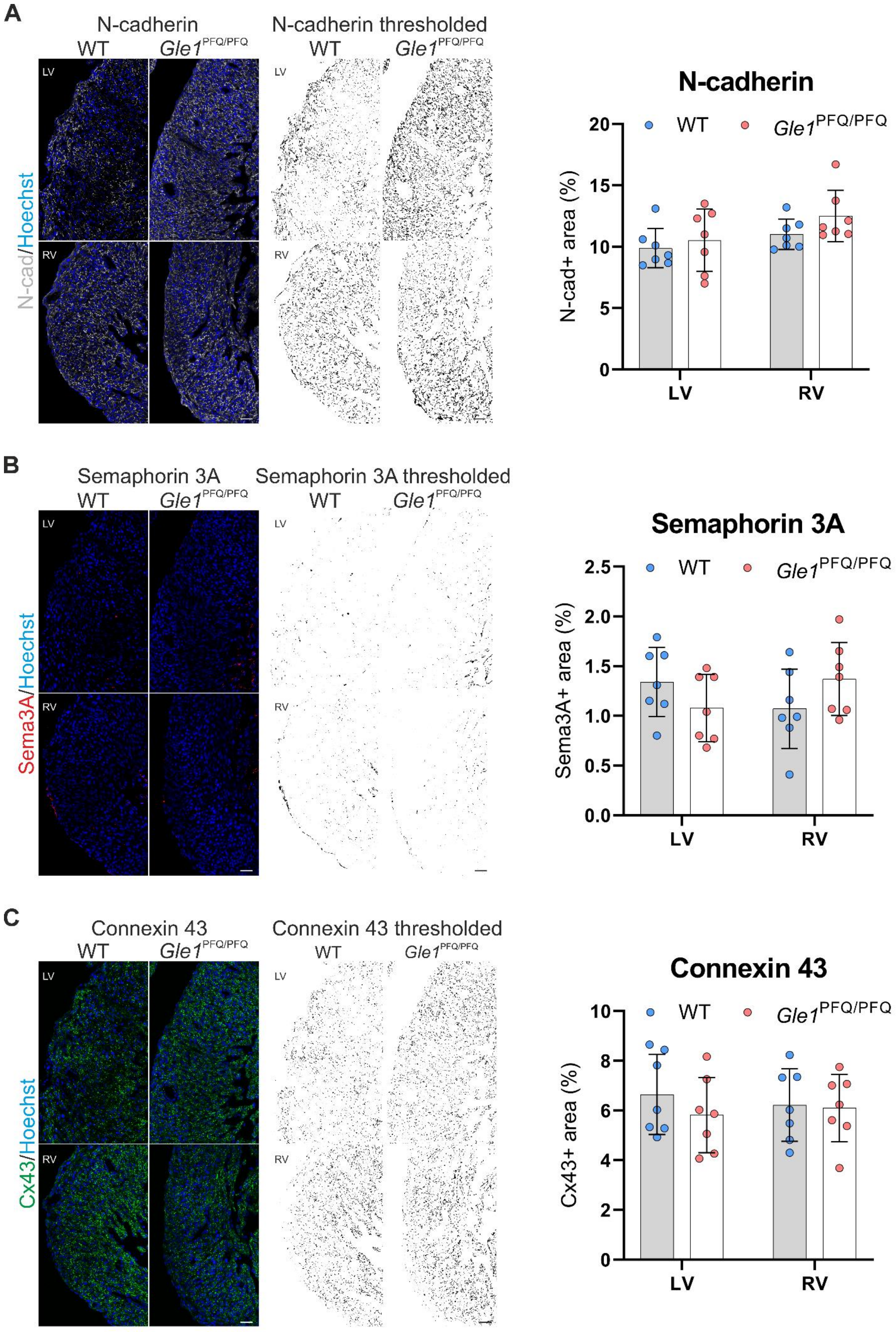


**Supplementary figure 9**: Intercellular communication analysis in E16.5 hearts

N-CADHERIN **A)**, SEMAPHORIN 3A **B)**, and CONNEXIN 43 **C)** immunofluorescent staining and signal thresholding (right) for ImageJ analysis. Fluorescent-positive area was expressed as percentage of total cross-sectional area of compact myocardium of the left (LV) and right (RV) ventricle in wildtype (WT) and *Gle1*$^{PFQ/PFQ}$ littermates. Means ± SD are plotted, as well as individual data points generated from each embryo (n=7 per genotype). Scale bar 40 µm.

**Supplementary figure 10**

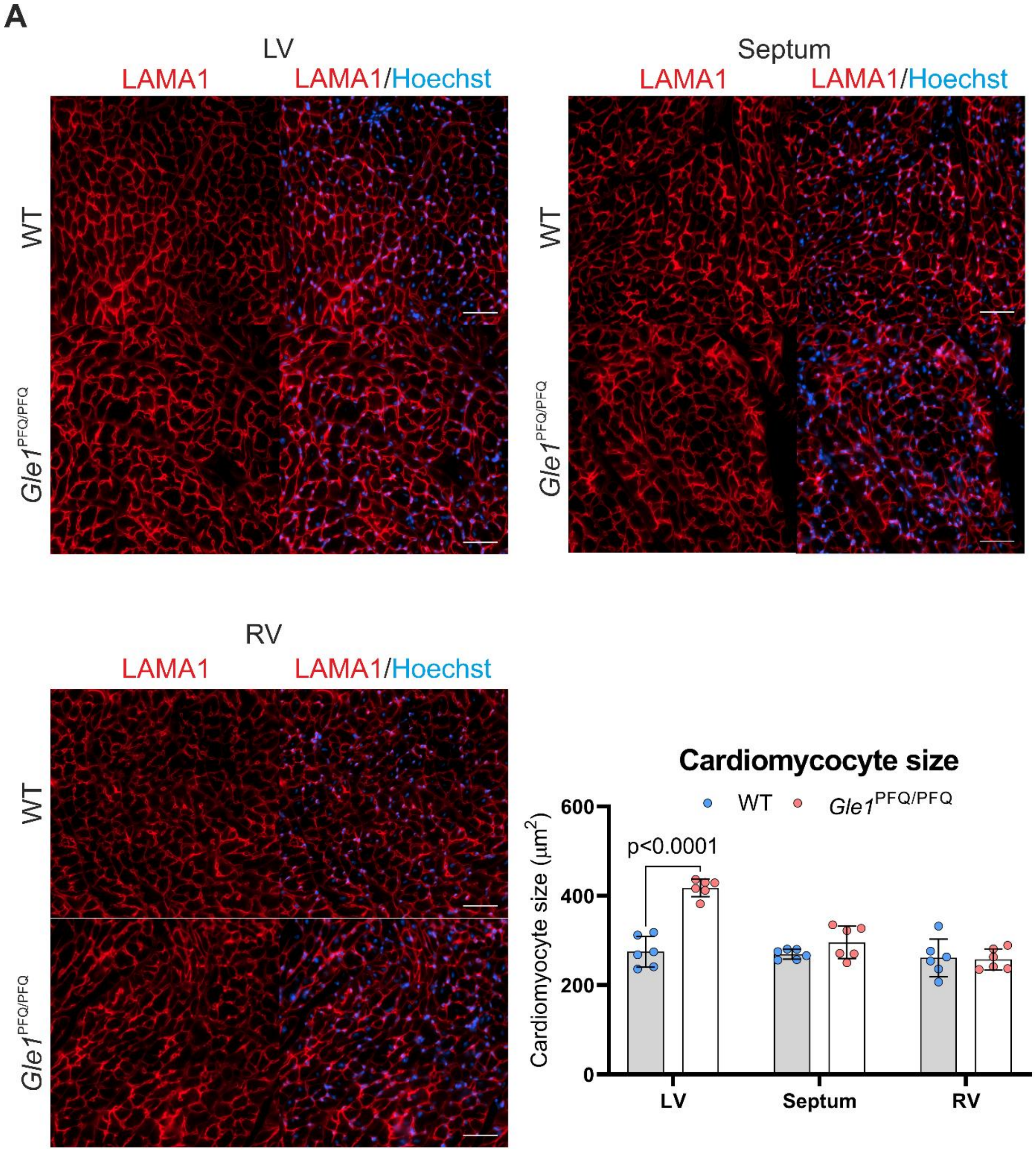


**Supplementary figure 10**: Analysis of adult heart cardiomyocytes

**A)** Laminin subunit α1 immunostaining of 25-week adult heart cross sections shown at the mid-level of the left ventricle (LV), septum, and right ventricle (RV). Quantification of cardiomyocyte size in each distinct heart region. Means ± SD are plotted, as well as individual data points generated from each mouse (over 100 cardiomyocytes in 3 fields per region, n=7 per genotype). Scale bar 50 µm, two-tailed Student's *t* test.

**Supplementary figure 11**

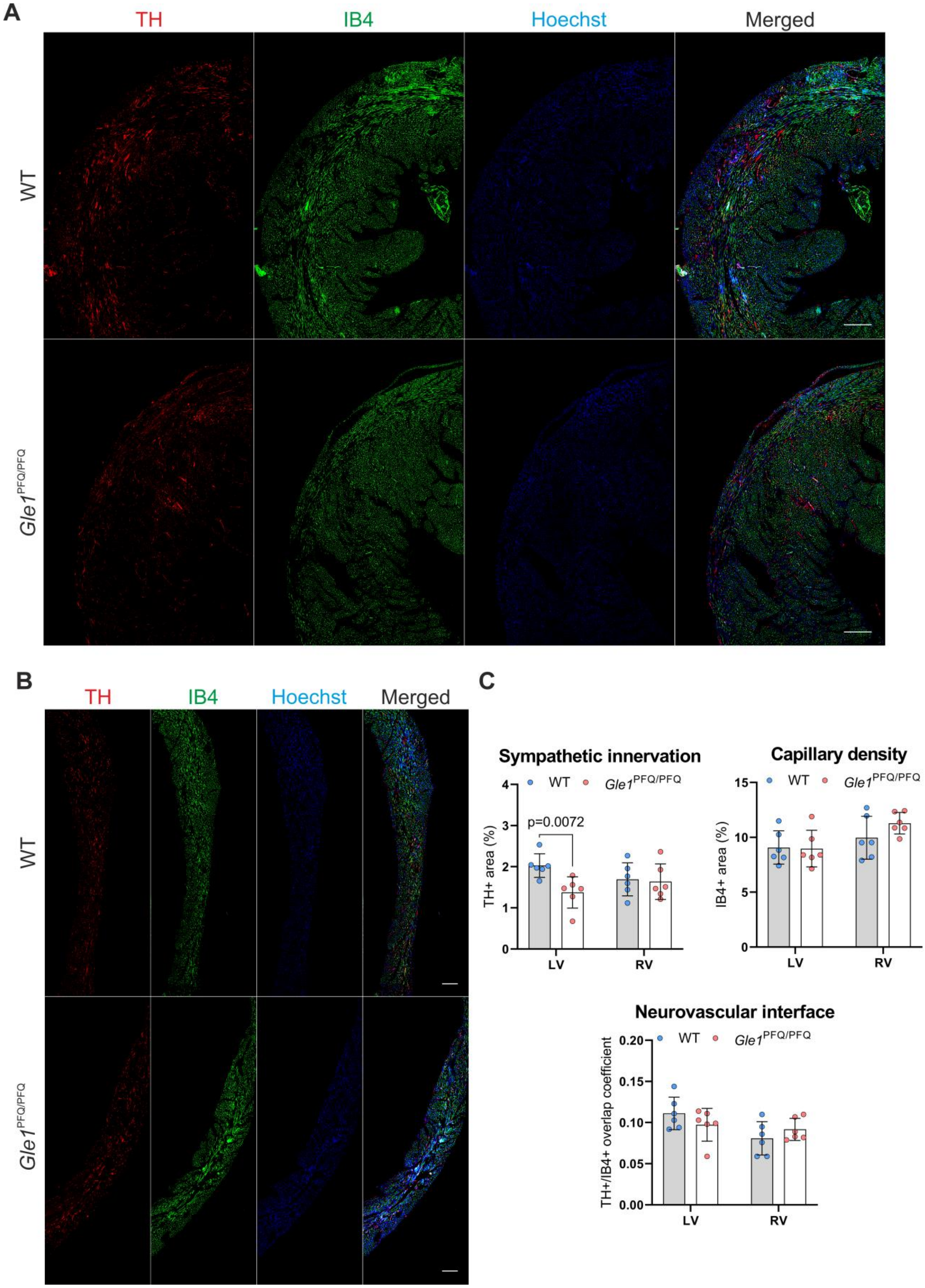

**Supplementary figure 11**: Characterization of adult heart sympathetic innervation and neurovascular interface

Representative images of 25-week wildtype (WT) and *Gle1*$^{PFQ/PFQ}$ adult heart immunostained for tyrosine hydroxylase (TH, red) and isolectine B4 (IB4, green) in **A)** left ventricles (LV) and **B)** right ventricle (RV). **C)** Quantification of TH-positive area in a single plane as a rate of sympathetic neuron innervation, IB4-positive area as a rate of vascularization, and TH vs IB4 colocalization as a rate of neurovascular interface. Means ± SD are plotted, as well as individual data points generated from mouse (n=7 per genotype). Scale bar 300 µm, two-tailed Student's *t* test

**Supplementary figure 12**

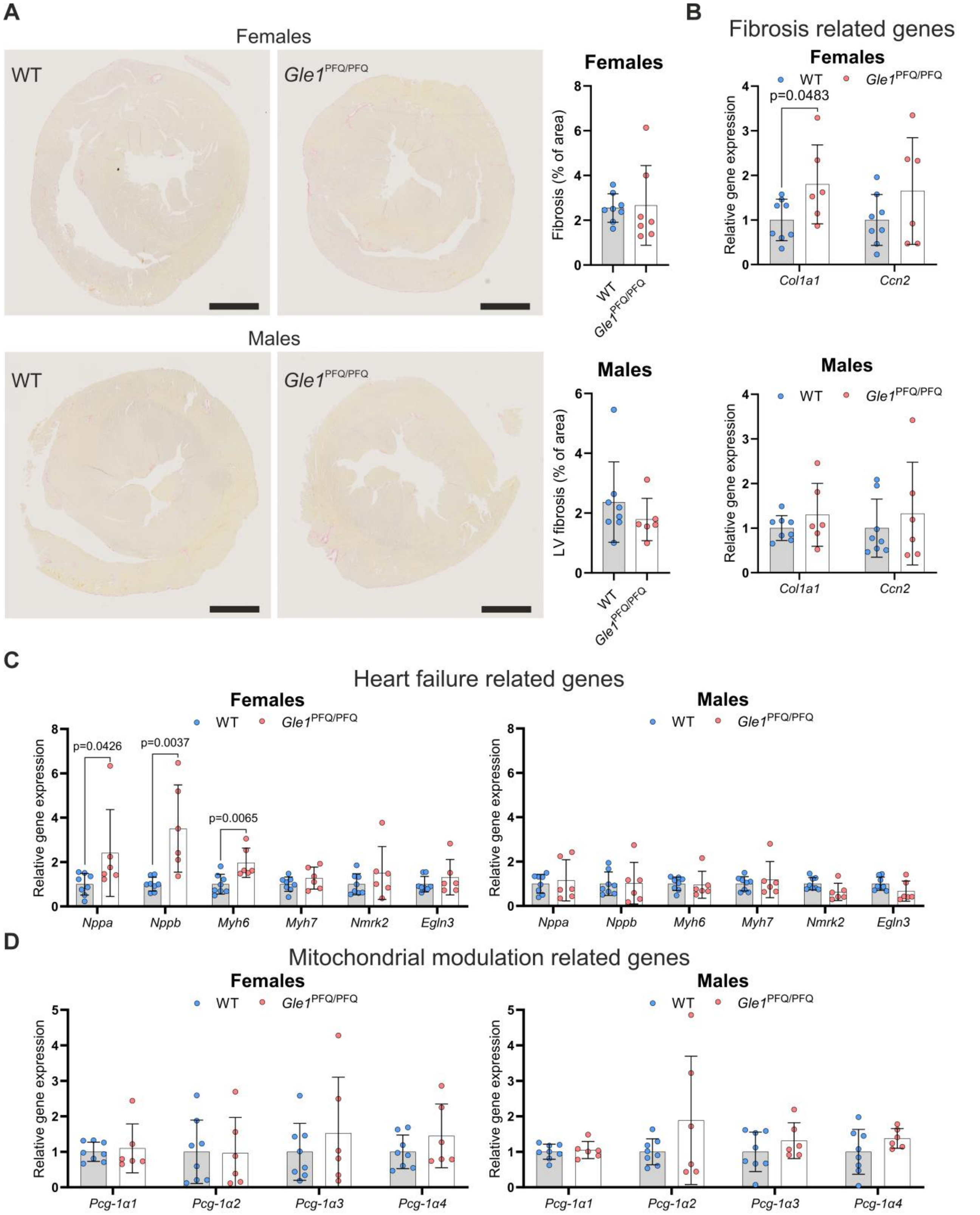


**Supplementary figure 12**: Molecular analysis of the *Gle1*$^{PFQ/PFQ}$ adult hearts

**A)** Adult heart (17-25 weeks) sections stained by picrosirius red to visualize the level of fibrosis. Means ± SD are plotted, as well as individual data points generated from each mouse (WT females: 8, KI females: 7, WT males: 8, KI males: 6). Scale bar 1 mm.

**B-D)** RT-qPCR analysis of adult hearts (17-25 weeks), focusing on transcripts related to **B)** fibrosis, **C)** heart failure, and **D)** mitochondrial modulation. Means ± SD are plotted, as well as individual data points generated from each mouse (WT females: 8, *Gle1*$^{PFQ/PFQ}$ females: 6, WT males: 8, *Gle1*$^{PFQ/PFQ}$ males: 6). Two-tailed Student's *t* test.

**Supplementary table 1**

Oligomers used for microinjections to modify the *Gle1* gene. The nine nucleotides inserted in the *Gle1* KI line via the ssDNA repair template are highlighted in red .

| | Oligo | Sequence | Manufacturer |
|---|---|---|---|
| KO line | crRNA1 | TGACATGCCATGCCGTCCGA | IDT |
| | crRNA2 | CTAGGGAGCCTGCTAAACGC | IDT |
| KI line | crRNA3 | CAGGTGCTGACCCTTCTAGG | IDT |
| | ssDNA | AGCAGATGAGCCAGGTGCTGTGCCACAGGGAAAACAGGTGCTGACCCTTCTAGCCGTTTCAAGAGGGCCTGAGGCAATGGCAGGAGGAGCAAGAGAGGAAGGTGCGAGCCCTCTCAGAGATGGC | IDT |

**Supplementary table 2**

Genotyping and sequencing primers for the Gle1 KO and KI mice

| Primer | Sequence | Manufacturer |
|---|---|---|
| *Gle1* KO DNA Se | CCACGTAGGCTCGCTAGTTT | IDT |
| *Gle1* KO DNA As | AGCTGAAGGCCCAAGCAAAT | IDT |
| *Gle1* KI DNA Se | ACAAGTCAGTGCTTGAATATAGAGC | IDT |
| *Gle1* KI DNA As | AACAGGCACTCAGGAAGACT | IDT |
| *Gle1* KI cDNA Se | GAGTCCCCGTTCACAGAACC | IDT |
| *Gle1* KI cDNA As | GAAGCTTTAGGTTGAGAATCTTGGC | IDT |

**Supplementary table 3**

List of RT-qPCR primers

| Primer | Sequence | Manufacturer |
|---|---|---|
| *Gle1* F | CTTCTTGGAGGTGTGCGGGA | IDT |
| *Gle1* R | TGTCCTGAGCTCGTGATGGC | IDT |
| *Cdkn2a* (p16) F | GTGTGCATGACGTGCGGG | IDT |
| *Cdkn2a* (p16) R | GCAGTTCGAATCTGCACCGTAG | IDT |
| *Cdkn2a* (p19) F | GCTCTGGCTTTCGTGAACAT | IDT |
| *Cdkn2a* (p19) R | TCGAATCTGCACCGTAGTTGAG | IDT |
| *Cdkn1a* (p21) F | AACATCTCAGGGCCGAAA | IDT |
| *Cdkn1a* (p21) R | TGCGCTTGGAGTGATAGAAA | IDT |
| *Il1α* F | GCACCTTACACCTACCAGAGT | IDT |
| *Il1α* R | AAACTTCTGCCTGACGAGCTT | IDT |
| *Il1β* F | TTCAGGCAGGCAGTATCACTC | IDT |
| *Il1β* R | GAAGGTCCACGGGAAAGACAC | IDT |
| *Il6* F | TAGTCCTTCCTACCCCAATTTCC | IDT |
| *Il6* R | TTGGTCCTTAGCCACTCCTTC | IDT |
| *Cxcl1* F | AACCGAAGTCATAGCCACAC | IDT |
| *Cxcl1* R | GACACCTTTTAGCATCTTTTGG | IDT |
| *Pai-1* F | TGGGTGGAAAGGCATACCAAA | IDT |
| *Pai-1* R | AAGTAGAGGGCATTCACCAGC | IDT |
| *Actb* F | CAGCAAGCAGGAGTACGATG | IDT |
| *Actb* R | GGTGTAAAACGCAGCTCAGT | IDT |
| *Ywhae* F | CCGAGCGATACGACGAAATG | IDT |
| *Ywhae* R | CAGTTTCAACCATTTGCCGG | IDT |
| 18s F | CGCCGCTAGAGGTGAAATTC | IDT |
| 18s R | CCAGTCGGCATCGTTTATGG | IDT |
| *Pgc-1α1* F | GGACATGTGCAGCCAAGACTCT | IDT |
| *Pgc-1α1* R | CACTTCAATCCACCCAGAAAGCT | IDT |
| *Pgc-1α2* F | CCACCAGAATGAGTGACATGGA | IDT |
| *Pgc-1α2* R | GTTCAGCAAGATCTGGGCAAA | IDT |
| *Pgc-1α3* F | AAGTGAGTAACCGGAGGCATTC | IDT |
| *Pgc-1α3* R | TTCAGGAAGATCTGGGCAAAGA | IDT |
| *Pgc-1α4* F | TCACACCAAACCCACAGAAA | IDT |
| *Pgc-1α4* R | CTGGAAGATATGGCACAT | IDT |
| *Nmrk2* F | GCCGTATGAGGAATGCAAGC | IDT |
| *Nmrk2* R | GCCATCTAAATAGACCACTTCCACC | IDT |

**Supplementary table 4**

List of TaqMan probes

| Primer | Forward | Reverse | Probe |
|---|---|---|---|
| 18s | TGGTTGCAAAGCTGAAACTTAAAG | AGTCAAATTAAGCCGCAGGC | CCTGGTGGTGCCCTTCCGTCA |
| *Nppa* | GAAAAGCAAACTGAGGGCTCTG | CCTACCCCCGAAGCAGCT | TCGCTGGCCCTCGGAGCCT |
| *Nppb* | TGGGCAGAAGATAGACCGGA | ACAACCTCAGCCCGTCACAG | CGGCGCAGTCAGTCGCTTGG |
| *Col1a1* | CCCTGGCCTTGGAGGAA | CACGGAAACTCCAGCTGATTTT | CTTTGCTTCCCAGATGTCCTATGGCTATGATG |
| *Ctgf* | CGCCAACCGCAAGATTG | CACGGACCCACCGAAGAC | CACTGCCAAAGATGGTGCACCCTG |
| *Myh6* | GGTGCCAAGAAGATGCACG | TTATGTTTATTGTGTATTGGCCACAG | CGAGGAATAACCTCTCCAGCAGACCCTC |
| *Myh7* | AGCTCTAAGGGTGCCCGTG | TGCTTCCACCTAAAGGGCTG | AGCCCTCAGACCTGGAGCCTTTGC |
| *Egln3* | TGAGGCTGGATCTGGAGAAGA | AGCCGACCTCGTGCAGAC | CGCCCTGGAGTACATCGTGCCC |

**Supplementary table 5**

List of antibodies used for western blot

| Target | Host | ID | Manufacturer | Dilution |
|---|---|---|---|---|
| GLE1 | Rabbit | ab96007 | Abcam | 1:500 |
| Anti-Puromycin | Rat | MABE341 | Merck | 1:25 000 |
| eIF2α | Rabbit | ab26197 | Abcam | 1:1 000 |
| eIF2α (phospho S51) | Rabbit | ab32157 | Abcam | 1:500 |
| Anti-rabbit-HRP | Sheep | A16172 | **Thermo Fisher** | 1:5 000 |
| Anti-rat-HRP | Goat | 31470 | **Thermo Fisher** | 1:50 000 |

**Supplementary table 6**

List of antibodies and probes used for fluorescent staining

| Target | Host | ID | Manufacturer | Dilution |
|---|---|---|---|---|
| GLE1 | Rabbit | ab96007 | Abcam | 1:500 |
| ZO-1 | Mouse | 339100 | Invitrogen | 1:400 |
| Tyrosine hydroxylase | Rabbit | 657012 | Merck | 1:500 (whole-mount) |
| Tyrosine hydroxylase | Mouse | MAB318 | Merck | 1:1000 (section) |
| SOX10 | Rabbit | AB155279 | Abcam | 1:250 |
| G3BP1 | Rabbit | 13057-2-AP | Thermo Fisher | 1:500 |
| HH3 (pSer10) | Rabbit | 06-570 | Thermo Fisher | 1:500 |
| γH2A.X (phospho S139) | Rabbit | ab81299 | Abcam | 1:200 |
| LAMA1 | Rabbit | L9393 | Merck | 1:1 000 |
| Connexin 43 | Rabbit | C6219 | Thermo Fisher | 1:1 000 |
| N-cadherin | Mouse | MA1-91128 | Thermo Fisher | 1:400 |
| Sema3A | Goat | sc-1148 | Santa Cruz | 1:50 |
| Vinculin | Mouse | V9131 | Merck | 1:500 |
| CDX2 | Mouse | CDX2-88 | BioGenex | 1:500 |
| Cleaved caspase-3 | Rabbit | 9664 | Cell Signalling | 1:600 |
| GATA6 | Goat | AF1700 | R&D Systems | 1:150 |
| NANOG | Rabbit | 8822 | Cell Signalling | 1:1 000 |
| OCT3/4 | Goat | sc-8628 | Santa Cruz | 1:400 |
| Isl1/2 | mouse | 39.4D5 | DSHB | 1:100 |
| Sox2 | goat | AF2018 | R&D Systems | 1:200 |
| Olig2 | rabbit | AB9610 | Merck | 1:500 |
| Alexa Fluor® 488 AffiniPure™ Donkey Anti-Goat IgG (H+L) | Donkey | 705-546-147 | Jackson ImmunoResearch | 1:400 |
| Donkey anti-Mouse IgG (H+L) Alexa Fluor™ 568 | Donkey | A10037 | Thermo Fisher | 1:400 |
| Donkey anti-Rabbit IgG (H+L) Alexa Fluor™ Plus 647 | Donkey | A32795 | Thermo Fisher | 1:400 |
| Alexa Fluor™ 488 Phalloidin | | A12379 | Thermo Fisher | 1:600 |
| Griffonia Simplicifolia Lectin I (GSL I) Isolectin B4. Biotinylated | | B-1205 | Vector laboratories | 1:50 |
| Streptavidin / FITC | | F0422 | DakoCytomation | 1:200 |
